# Spatial phylogenetics of Japanese ferns: Patterns, processes, and implications for conservation

**DOI:** 10.1101/2021.08.26.457744

**Authors:** Joel H. Nitta, Brent D. Mishler, Wataru Iwasaki, Atsushi Ebihara

## Abstract

**Premise:** Biodiversity is often only measured with species richness. However, this ignores evolutionary history and is not sufficient for making conservation decisions. Here, we characterize multiple facets and drivers of biodiversity to understand how these relate to bioregions and conservation status in the ferns of Japan.

**Methods:** We compiled a community dataset of 1,239 20 km × 20 km grid-cells including 672 taxa based on > 300,000 specimen records. We combined this with a phylogeny and functional traits to analyze taxonomic, phylogenetic, and functional diversity, and modeled biodiversity metrics in response to environmental factors and reproductive mode. Hierarchical clustering was used to delimit bioregions. Conservation status and threats were assessed by comparing the overlap of significantly diverse grid-cells with conservation zones and range maps of native Japanese deer.

**Results:** Taxonomic richness was highest at mid-latitudes. Phylogenetic and functional diversity and phylogenetic endemism were highest in small southern islands. Relative phylogenetic and functional diversity were high at high and low latitudes, and low at mid-latitudes. Grid-cells were grouped into three (phylogenetic) or four (taxonomic) major bioregions. Temperature and apomixis were identified as drivers of biodiversity patterns. Conservation status was generally high for grid-cells with significantly high biodiversity, but the threat due to herbivory by deer was greater for taxonomic richness than other metrics.

**Conclusions:** Our integrative approach reveals previously undetected patterns and drivers of biodiversity in the ferns of Japan. Future conservation efforts should recognize that threats can vary by biodiversity metric and consider multiple multiple metrics when establishing conservation priorities.

---

Characterizing the spatial distribution of biodiversity is a major goal of evolutionary biology with two equally important aspects: to understand the processes generating it, and to conserve it. Until recently, the vast majority of studies seeking to understand the spatial distribution of biodiversity have focused on species richness. Hotspots, or areas of exceptionally high species richness or endemism, have received particular attention, resulting in a widely recognized set of 36 terrestrial hotspots (Myers et al., 2000; Noss, 2016) and motivating conservation strategies to preserve them (Margules and Pressey, 2000; Brooks et al., 2006). However, species richness alone cannot provide a complete picture of biodiversity (Miller et al., 2018). All organisms are related to a greater or lesser degree by descent from a common ancestor, and these evolutionary relationships must be taken into account to obtain a full understanding of the biodiversity present in an area.

Presumably, the main reason phylogeny has not been taken into account more prominently in biodiversity studies is because the necessary data (DNA sequences and georeferenced occurrence records) and analytic tools have only become available relatively recently (Soltis and Soltis, 2016; Folk and Siniscalchi, 2021). These datasets and tools now make it possible to analyze other dimensions of biodiversity, such as phylogenetic diversity (Faith, 1992) and phylogenetic endemism (Rosauer et al., 2009). Better ways to characterize biological regions (“bioregions”; *ie*, areas defined by their taxonomic composition or evolutionary history) are also available (Laffan et al., 2016; White et al., 2019; Daru, Karunarathne, et al., 2020), rather than relying on ad-hoc characterizations. Furthermore, incorporating these two frameworks—the categorization of areas into phyloregions with analysis of over/under dispersion of biodiversity—can provide powerful insights into the processes structuring biodiversity and suggest conservation priorities.

However, a comprehensive understanding of the relationships between richness and other metrics requires densely sampled data, and such datasets are rare on the regional (country) scale. The ferns of Japan are excellent model system because they have been the target of intense botanical interest for several decades and are densely sampled (reviewed in Ebihara and Nitta, 2019). The ferns of Japan include 676 native, non-hybrid taxa (including species and varieties) and hundreds of hybrids (nothotaxa; Ebihara and Nitta, 2019). The availability of detailed distribution data (distribution maps at the ca. 10 km scale for nearly all species; Ebihara, 2016, 2017), trait data (multiple quantitative and qualitative traits compiled for identification of nearly all species; Ebihara and Nitta, 2019), and DNA sequences for > 97% of species (Ebihara et al., 2010; Ebihara and Nitta, 2019; all coverage statistics exclude nothotaxa) make the ferns of Japan an ideal system for investigating the relationships between, and drivers of, multiple dimensions of biodiversity.

One particularly valuable characteristic of the Japanese fern flora is the availability of data on reproductive mode, which is known for 492 native fern taxa excluding hybrids (72.8%; Ebihara and Nitta, 2019). Reproductive mode is likely to affect population-level genetic diversity (Bengtsson, 2003), and thereby higher-level biodiversity (Krueger-Hadfield et al., 2019). In ferns, clonally reproducing apomictic species are often polyploid hybrids that share identical plastid genotypes among taxa and within populations (Grusz et al., 2009; Chao et al., 2012; Hori et al., 2014; Hori and Murakami, 2019). Therefore, we expect high prevalence of apomictic species within a community to decrease phylogenetic diversity.

Japan is situated along a latitudinal gradient spanning seasonal, temperate areas in the north to subtropical, mostly aseasonal islands in the south. Furthermore, it is a mountainous country with great variation in elevation on the larger islands. Plant distributions often reflect physiological adaptations to climate (*eg*, precipitation and temperature; Woodward, 1987). Therefore, we expect the spatial distribution of biodiversity to be determined by climatic variation in addition to reproductive mode.

Here we leverage this exceptionally rich dataset to analyze the geographic distribution of biodiversity in detail. We ask the following questions in our study system, the ferns of Japan: 1. How is biodiversity distributed? 2. How is biodiversity structured? 3. What environmental and biological factors influence biodiversity? 4. How well is biodiversity protected?

## MATERIALS AND METHODS

All computational analyses were carried out in R v4.1.2 (R Core Team, 2021) unless otherwise stated. The R package ‘targets’ v.0.9.0 was used to control analysis workflow (Landau, 2021).

### Study Site

Japan (20°N to 45°N, 122°E to 153°E) consists of four main islands and thousands of smaller ones (Fig. 1). Most of the main islands have been in contact with the continent at various points in the past during periods of lower sea level, but the Ogasawara archipelago consists of oceanic islands that have never been connected to the mainland. The climate is subtropical in the southern islands and temperate in other parts of the country. Map data for Japan were downloaded from the Geospatial Information Authority of Japan under the Creative Commons Attribution License v4.0 (https://www.gsi.go.jp/kankyochiri/gm_japan_e.html).

**Fig 1.**
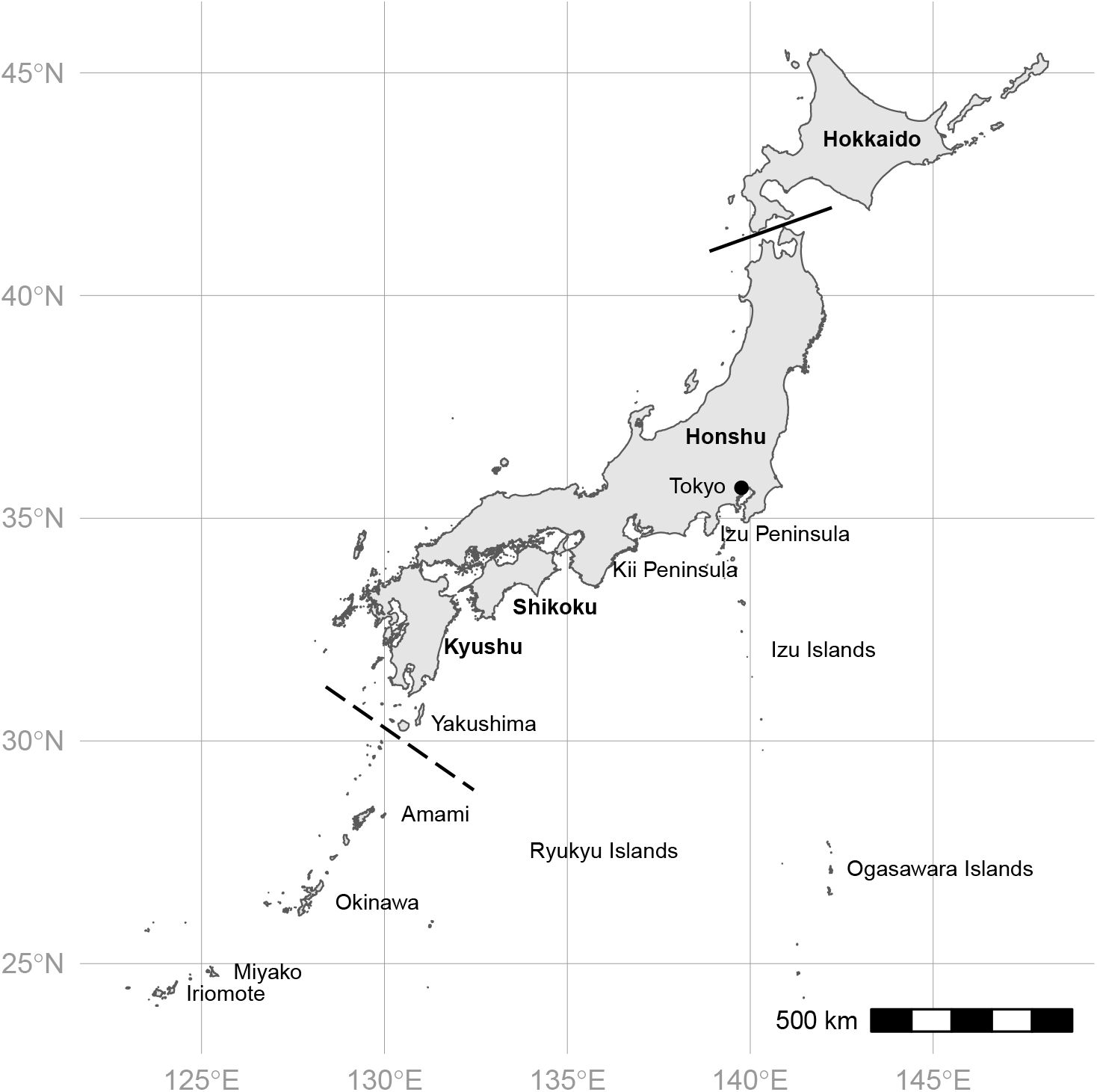
Map of Japan showing names of places mentioned in the text. Names of the four main islands in bold. The Ryukyu Islands include Yakushima, Amami, Okinawa, Miyako, and Iriomote. Solid line shows location of Tsugaru Strait; dashed line shows Tokara Strait.

### Occurrence data

We used a list of native, non-hybrid fern specimens deposited at herbaria in Japan to quantify occurrences (Ebihara, 2016, 2017; Ebihara and Nitta, 2019). We chose to use this over other sources such as the Global Biodiversity Information Facility (GBIF) because of its high quality, which obviates many cleaning steps otherwise needed when working with large, publicly available datasets. Furthermore, the overall sampling completeness of this dataset is also high (Fig. S1 in Appendix S1; see Supplemental Data with this article). The original list comprises 337,000 specimens representing 673 terminal taxa at the rank of species, subspecies, form, or variety, excluding non-native taxa and nothotaxa. We treated infraspecific taxa as distinct species during analysis and hereafter use “taxon” and “species” interchangeably to refer to entities at this rank, unless otherwise indicated. All names are standardized to a common taxonomy, the Japan Green List v. 1.01 (http://www.rdplants.org/gl/).

Occurrence records were georeferenced and thoroughly vetted as follows (Ebihara, 2016, 2017; Ebihara and Nitta, 2019). For specimens that lacked latitude and longitude data, georeferencing was done by mapping collection site names to a set of standard ca. 10 km × 10 km grid squares defined by the Statistics Bureau of Japan (the “second-degree mesh”; http://www.stat.go.jp/english/data/mesh/05.html), and the centroid of the second-degree mesh cell used as the specimen location. In the case that the collection site could not be mapped to a single second-degree mesh cell, it was excluded. Next, occurrence maps were generated for each taxon showing presence or absence in each second-degree mesh cell. In the case that a given taxon appeared insufficiently sampled (*ie*, not present on the map where it would typically be expected to occur), AE and members of the Nippon Fernist Club (local botanists familiar with the ferns of Japan) searched for additional specimens, which were then added to the list. The occurrence maps were iteratively refined until the vast majority of known Japanese ferns had been observed across their expected ranges; the resulting set of occurrence records can be considered accurate to ca. 10 km (the grain size of the second-degree mesh map).

We further cleaned the list prior to analysis by filtering out any occurrences not within the second-degree mesh and removing duplicate collections (300,685 specimens, 673 taxa after filtering). Given the high quality of our occurrence data and the fact that automated occurrence cleaning algorithms (*eg*, CoordinateCleaner; Zizka et al., 2019) have the potential to erroneously exclude true occurrence points (*ie*, false positives; Zizka et al., 2020), we chose not to apply additional automated cleaning steps to our data as is often done with occurrence records obtained from GBIF (*eg*, Rice et al., 2019; Suissa et al., 2021).

A necessary step in any analysis of biodiversity is to set the grain size used to accurately define communities (co-occurring species). Here, one must consider that smaller grain size is needed to detect environmental effects, while larger grain size is needed to ensure adequate species sampling. We created sets of 10 km, 20 km, 30 km, and 40 km grid-cells covering Japan using a Mollweide equal-area projection (size refers to the length of the side of each square grid-cell). At each grain size, species occurrences were converted to a presence-absence community matrix (a species was considered present if at least one specimen was recorded in that grid-cell). We calculated sampling redundancy (1 - richness/number of specimens; Garcillán et al., 2003) to quantify adequacy of sampling. Preliminary analysis indicated that 20 km grid-cells are optimal for our dataset: there is a sudden improvement in redundancy values from 10 km to 20 km, but much less improvement as grain size is increased beyond 20 km (Fig. S2). Although the grid-cells are defined with equal area, actual land area of each cell varies due to coastlines. Strictly filtering out all grid-cells with less than complete land area would result in a large loss of data, as Japan has many coastlines and small islands. Therefore, we instead filtered the grid-cells by sampling completeness, removing any cells with redundancy < 0.1; this dataset was used for all subsequent analyses. The R package ‘sf’ v.1.0.5 was used for all GIS analyses (Pebesma, 2018).

### Morphological trait data

We used traits originally compiled for identification of ferns and lycophytes of Japan (Ebihara, 2016, 2017), which were formatted to be used with Lucid software (https://www.lucidcentral.org/). Continuous traits were measured on 10 randomly selected specimens per species, then four values per species were obtained from these measurements following Lucid format: outside (*ie* outside the typical range) minimum, typical minimum, typical maximum, and outside maximum. For species with dimorphic fertile (*ie*, spore-bearing) and sterile fronds, fertile and sterile fronds were measured separately. We took the mean of the typical minimum and maximum values to use as the species mean value. Qualitative traits were scored by observing voucher specimens. All qualitative traits were scored in binary format; *eg*, a single trait with three states was formatted as three binary traits. From the original trait list, we selected only traits with putative ecological function (Table 1) and excluded any traits that had Pearson correlation coefficient > 0.6 or fewer than three observations of a given trait state.

**Table 1.** Fern traits used in this study. Letters in parentheses indicate trait type: b, binary (trait present or absent); c, continuous; q, qualitative. Qualitative traits were coded as binary for analysis (see Materials and Methods). For complete list of trait states for qualitative traits, see Table S3.

| Trait | Significance | Reference |
| --- | --- | --- |
| Indusium (b) | Protects developing spores | Poppinga et al. (2015) |
| Frond width (c) | Scales with overall plant size | Creese et al. (2011) |
| No. pinna pairs (c) | Less divided fronds have lower rates of evapotranspiration | Kluge and Kessler (2007) |
| Stipe length (c) | Shorter lengths minimize resistance | Watkins Jr. et al. (2010) |
| Frond margin (q) | Thermoregulation | Nicotra et al. (2011) |
| Frond shape (q) | Less divided fronds have lower rates of evapotranspiration | Kluge and Kessler (2007) |
| Frond texture (q) | Thicker leaves have lower rates of evapotranspiration | Givnish (1987) |

### Reproductive mode data

We used data compiled by Ebihara and Nitta (2019), which classifies each fern taxon as sexual, apomictic, mixed (both sexual and apomictic modes known from the same taxon), or unknown. We calculated the percentage of apomictic taxa in each grid-cell as the sum of apomictic and mixed taxa divided by total number of taxa (Fig. S3B).

### Environmental data

We downloaded climate data at the 2.5 minute scale from the WorldClim database using the ‘getData’ function in the R package ‘raster’ v.3.5.11 (Hijmans, 2021), and extracted four variables related to our research questions as follows: mean annual temperature (bio1; hereafter “temperature”), temperature seasonality (bio4), annual precipitation (bio12; hereafter “precipitation”), and precipitation seasonality (bio15) (Fig. S3). We identified the intersection of the climate data at the 2.5 minute scale with the 20 km grid-cells, then calculated the mean of each climatic variable for each grid-cell. We calculated area as rolling mean area (km^2^) across 100 km latitudinal bands. This approach reflects the idea that the amount of area with a given climate regime enclosing each grid-cell may influence species richness of that cell (Fahrig, 2013), and that using raw island areas would not be appropriate in this context as there are just four very large islands with many much smaller ones located close by (Fig. 1).

### Phylogenetic analysis

Sequences for the plastid, coding *rbcL* gene are available for ca. 97% of the Japanese fern flora (Ebihara and Nitta, 2019); these sequences originate from samples that were identified using the same taxonomic system as the occurrence data (Ebihara et al., 2010; Ebihara and Nitta, 2019), so we used these preferentially over other data available on GenBank that might include misidentifications or different taxon concepts. However, using this dataset alone for phylogenetic analysis suffers two drawbacks: it is not densely sampled enough to resolve some internal nodes, and it lacks many lineages that have fossils available for molecular dating. Furthermore, use of community phylogenies generated by only by sampling the species present in the local community has been shown to produce spurious results in simulation studies (Park et al., 2018). Because one goal of the study is to understand the distribution of paleo-vs. neo-endemic areas (defined in units of time), we require an ultrametric tree. Therefore, to obtain a robust, ultrametric tree, we combined the *rbcL* data of Ebihara and Nitta (2019) (706 taxa) with a globally sampled plastid dataset including all fern sequences on GenBank for four widely-sequenced plastid genes (*atpA, atpB, rbcL*, and *rps4*; 4,492 taxa) and 58 other coding, single-copy plastid genes extracted from 123 complete fern plastomes (Nitta et al., in prep.) (4,787 OTUs total, including outgroup taxa). This gene sampling is comparable to a recent global fern phylogeny that resolved relationships across ca. 4,000 taxa using six plastid markers (Testo and Sundue, 2016), and the addition of plastome data can be expected to increase support along the backbone.

We conducted maximum-likelihood phylogenetic analysis with IQ-TREE v1.6.12 (Nguyen et al., 2015) under the GTR+I+G model (all sites concatenated into a single data matrix), and assessed node support with 1,000 ultrafast bootstrap replicates (Minh et al., 2013). Molecular dating was conducted with treePL v1.0 (Smith and O’Meara, 2012) using 26 fossil calibration points after Testo and Sundue (2016), with the exception of treating fossil *Kuylisporites* as belonging to crown *Cyathea*, not *Cyathea* + *Alsophila* (Loiseau et al., 2020). The root of the tree (most recent common ancestor of all land plants) was fixed to 475 mya (Testo and Sundue, 2016). We trimmed the resulting ultrametric tree to only Japanese taxa. treePL requires branch lengths to be > 0, and it sets extremely small branches to an arbitrary minimum positive length. This may artificially add branch length between taxa with identical DNA sequences, which occur multiple times in the Japanese fern *rbcL* dataset. We converted any clades in the dated tree consisting of identical sequences to polytomies with the node at 0 mya (*ie*, the present). This tree was used for all subsequent analysis unless indicated otherwise.

### Phylogenetic signal

We tested for phylogenetic signal in continuous traits using two alternative metrics, Blomberg’s *K* (Blomberg et al., 2003) and Pagel’s lambda (Pagel, 1999). *K* describes the ratio between the observed variance in a trait vs. the amount of variance expected under Brownian Motion (BM); lambda is a scaling parameter between 0 and 1 that transforms the phylogeny such that trait values fit those most closely expected under BM. Both lambda and *K* = 1 under BM; relatively lower values indicate less signal (overdispersion) and higher values indicate more signal (clustering). We measured *K* and lambda using the ‘phylosig’ function in the R package ‘phytools’ v.0.7.90 (Revell, 2012), which also calculates significance based on a randomization test for *K* and a likelihood ratio test for lambda. We tested for phylogenetic signal in qualitative (coded as binary) traits using Fritz and Purvis’ *D* (Fritz and Purvis, 2010) with the ‘phylo.d’ function in the R package ‘caper’ v.1.0.1 (Orme et al., 2018), which also assesses significance by comparing the observed value to a null distribution of random values and a simulated distribution of values expected under BM. Values of *D* range from 0 under BM to 1 under random evolution, but can exceed this range in cases of extreme clumping or overdispersion, respectively. These methods for phylogenetic signal cannot use a tree with zero-length polytomies, so we used the original dated tree from treePL.

### Analysis of biodiversity

We measured taxonomic diversity using taxonomic richness (number of taxa in each community).

We measured phylogenetic diversity (PD) using the Faith’s PD (the total branch length connecting all terminal taxa in a community, including the root of the phylogeny; Faith, 1992). To detect areas with unusually long or short branches, we calculated relative phylogenetic diversity (RPD), which is a ratio of observed PD to PD measured on a tree where all branch lengths have been transformed to equal length (Mishler et al., 2014).

We measured functional diversity (FD) following the method of Petchey and Gaston (2002). We first calculated functional distances using Gower’s method (Gower, 1971) on the trait data with the ‘gowdis’ function in the R package ‘FD’ v.1.0.12 (Laliberté and Legendre, 2010). We weighted continuous and qualitative traits equally, weighted each binary component trait of each qualitative trait equally, and log-transformed and scaled continuous traits prior to calculating distances. We also tested alternative weighting schemes, but these produced very similar results (data not shown). We then used the distances to build a trait dendrogram using the ‘hclust’ function in the R package ‘stats’ v.4.1.2 (R Core Team, 2021), and used the dendogram in place of a phylogenetic tree to calculate PD and RPD as described above. These functional analogs of PD and RPD are hereafter referred to as FD (functional diversity) and RFD (relative functional diversity).

We measured phylogenetic endemism (PE) as the sum of all branches at a site inversely weighted by their range size (Rosauer et al., 2009).

Observed values of PD, RPD, FD, RFD, and PE are known to be related to species richness: as more taxa are drawn at random without replacement from a given set, it becomes increasingly unlikely to draw a taxon that is distantly related (or functionally dissimilar) to the others.

Therefore, we determined whether the observed metrics are more or less diverse than random for a given number of taxa by conducting a randomization test. We generated 999 random communities using the “curveball” algorithm of Strona et al. (2014), which conserves richness per site and species occurrence frequencies while randomizing species identities, then compared the observed value to the distribution of random values. Statistical significance was assessed using a one-tailed test for PE or a two-tailed test for other metrics with alpha = 0.5; observed values in the extreme 5% of the null distribution were considered significant. We also calculated the standard effect size (SES) of each biodiversity metric. SES measures how extreme the observed value (*obs*) is relative to the null distribution (*null*):

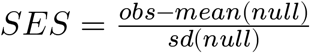

We tested whether centers of endemism have an over-representation of old (paleoendemic) or new (neoendemic) lineages using categorical analysis of neo- and paleo-endemism (CANAPE), which involves measuring PE with an alternate tree where all branch lengths are equal, then comparing the ratio between raw PE and alternative PE (RPE) (Mishler et al., 2014). Low RPE indicates short range-restricted branches (neoendemism) and high RPE indicates long range-restricted branches (paleoendemism). Since all of these measures are affected by species richness, categorization of endemism is done using the rank of each observed variable relative to those calculated for the null distribution. All measures of biodiversity and CANAPE were carried out using the R package ‘canaper’ v.0.0.2 (Nitta, 2022).

It is possible that endemism may be affected by species that are actually wide-ranging, but only have a small range in Japan (a “border effect”). This is especially expected for southern, sub-tropical islands, which harbor many tropical species with small ranges in Japan but wider ranges outside of the country. To partially account for this, we also calculated all endemism-related metrics using a dataset including species found only in Japan.

### Spatial models

Two grid cells lacking environmental data and an outlier with 100% apomictic taxa due to extremely low species richness were excluded from modeling analyses (Appendix S2; see Supplemental Data with this article).

We tested for spatial autocorrelation in biodiversity metrics (richness, SES of PD, SES of FD, SES of RPD, SES of RFD) and independent variables (environmental variables and % apomictic taxa) using Moran’s I with the ‘moran.mc’ function in the R package ‘spdep’ v.1.1.13 (Bivand et al., 2013). In all cases, significant spatial autocorrelation was detected (Moran’s I > 0, *P* value of permutation test < 0.05) (Table S1). Therefore, we used spatially explicit methods for modeling.

We checked for correlation between independent variables while taking into account spatial autocorrelation using a modified *t*-test with the ‘modified.ttest’ function in the R package ‘SpatialPack’ v.0.3.8196 (Vallejos et al., 2020). This indicated that temperature seasonality and % apomictic taxa were correlated with mean annual temperature (Table S2), so for the main modeling analysis we only included the following environmental variables: mean annual temperature, precipitation, and precipitation seasonality.

We constructed a linear mixed model for each biodiversity metric dependent on the environmental variables (“environmental models”). Initial inspection of the relationship between each biodiversity metric and temperature showed a hump-shaped pattern (Appendix S2), so we also included a quadratic term for temperature only. Richness was fit with the negative binomial response family; others (SES of PD, SES of FD, SES of RPD, SES of RFD) were Gaussian. We fit models including a Matérn correlation matrix based on the geographic centroids of each grid-cell as a random effect with the R package ‘spaMM’ v.3.9.25 (Rousset and Ferdy, 2014). We verified the significance of each term of the model by conducting likelihood ratio tests (LRTs) between the full model and models each with a single term of interest removed, in sequence. We also verified the significance of the full model relative to the null model (only including the Matérn correlation matrix) using LRTs.

We postulated that % apomictic taxa might explain the distribution of phylogenetic diversity better than temperature, so we also constructed a limited set of models to test this hypothesis (“reproductive models”). These models included biodiversity metrics with a phylogenetic component (SES of PD and SES of RPD) as dependent on % apomictic taxa, precipitation, and precipitation seasonality, with random spatial effects as described above. We compared the fit between environmental models and reproductive models for SES of PD and SES of RPD using conditional Akaike Information Criterion (cAIC) (Vaida and Blanchard, 2005; Courtiol and Rousset, 2017).

### Analysis of bioregions

We analyzed bioregions using the framework of Daru, Farooq, et al. (2020) and their R package ‘phyloregion’ v.1.0.6. Briefly, this involves first calculating beta diversity (change in composition between communities) using either species names (taxonomic bioregions) or phylogenetic distances (phylogenetic bioregions). For taxonomic bioregions, we calculated the turnover in species between sites due to species replacement (Baselga, 2012) with the Sørensen index. For phylogenetic bioregions, we calculated phylogenetic dissimilarities based on the PhyloSor index (Bryant et al., 2008). The distances are used to construct a dendrogram, which is then split into into *k* bioregions. The optimal value of *k* cannot be known *a-priori*. Previous vegetation zone schemes in Japan have typically included 5–6 zones (Shimizu, 2014); to enable approximate comparison with these, we tested values of *k* from 1 to 10, and selected the optimal value using the ‘optimal_phyloregion’ function in the R package ‘phyloregion’ v.1.0.6.

### Assessment of conservation status and threats

We downloaded SHP files of conservation zones in Japan from the Japan Ministry of the Environment (https://www.biodic.go.jp/biodiversity/activity/policy/map/map17/index.html) and Ministry of Land, Infrastructure, Transport and Tourism (https://nlftp.mlit.go.jp/ksj/gml/datalist/KsjTmplt-A45.html). We excluded marine zones and those that do not protect plants. We categorized protected areas as either “high” (no human activities allowed at all) or “medium” status (some economic activities allowed by permit; Kusumoto et al., 2017); areas not afforded at least medium level of protection were not considered. Conservation status in Japan is administered by multiple laws, and protected areas frequently overlap (Natori et al., 2012). To prevent double-counting, all protected areas within a protection status were merged, and protected areas that overlapped between medium and high status were considered only high status.

Besides habitat loss due to human activity, one major threat to ferns in Japan is herbivory by Japanese deer (*Cervus nippon*). Although native, Japanese deer have caused extensive damage to native plant communities due to rapid population growth and range expansion since the 1970s (Takatsuki, 2009). Decrease or extirpation of fern populations due to deer herbivory has been observed in multiple sites and species across the country (Minamitani, 2005; Yahara, 2006; Hattori et al., 2010). Furthermore, 33 of 212 (15.5%) fern taxa listed as endangered (Red List status CR, EN or VU) in Japan include herbivory by deer as one of the reasons for their endangered status (Japan Ministry of the Environment, 2015, 2022). Although not all species of ferns are equally susceptible to deer herbivory (some are unpalatable to deer; Minamitani, 2005), these studies show it is an existential threat to many species. Therefore, analysis of the threat posed to fern biodiversity by deer herbivory is needed to inform future conservation policies.

We downloaded distribution maps (SHP and TIF) of *Cervus nippon* from the Japan Ministry of the Environment (https://www.biodic.go.jp/biodiversity/activity/policy/map/map14/index.html). To assess change in threat due to herbivory by deer over time, we used three maps available in this dataset: range of Japanese deer surveyed in 1978, range surveyed in 2003, and estimated range inferred from a model including snow cover and forest type based on the 2003 survey data (Japan Ministry of the Environment, 2021b). We limited estimated range to areas considered “highly likely” to be occupied by deer (movement cost < 0.10; Japan Ministry of the Environment, 2021b).

We cropped grid-cells to only include land area, then calculated the percent of protected area of each status type or percent of area occupied by deer for each survey period within grid-cells with high levels of biodiversity as measured with PD, FD, PE, or taxonomic richness. These values were compared with the overall percent of protected area or area occupied by deer across Japan (baseline rate). For PD, FD, and PE, significance was assessed with a one-tailed test at alpha = 0.05 against the 999 null communities described above. For taxonomic richness, cells ranked in the top > 5% were considered highly diverse.

## RESULTS

### Datasets

The phylogeny included 663 taxa (98.08% of the total number of native, non-hybrid ferns [676 taxa]) representing 98 genera and 34 families. The topology was in general agreement with other recent, large plastid fern phylogenies (Schuettpelz and Pryer, 2007; Lehtonen, 2011; Testo and Sundue, 2016; Fig. S4). Ferns were recovered as monophyletic, as were all families and genera except for *Microsorum* (families follow Pteridophyte Phylogeny Group I, 2016). The age of the fern clade is estimated to be 433.1 ma.

The cleaned list of specimens included 300,685 specimens representing 673 taxa (99.56%). The distribution of collection locations was highly skewed, with > 3,000 specimens per grid-cell northwest of Tokyo in Saitama prefecture, and many fewer elsewhere (Fig. S5A). Despite this, sampling redundancy was generally good (> 0.3; González-Orozco et al., 2014) throughout the country (mean 0.55 ± 0.17, final community data matrix; errors are SD unless mentioned otherwise; Fig. S5B). The final community data matrix included 672 taxa and 1,239 grid-cells. 138 grid-cells were excluded due to low redundancy (< 0.1); these were mostly from Hokkaido and coastal areas.

After removing correlated and low-variance traits, the final trait set included one binary trait, three continuous traits, and three qualitative traits (coded as 73 binary traits; Tables 1, S3. The trait matrix included 675 taxa (99.9%) and 77 traits, with 0.42% missing data.

### Phylogenetic signal

All continuous traits showed significant phylogenetic signal, but estimates of signal strength differed between *K* and lambda: all lambda values were > 0.95, indicating evolution by BM; however, values of *K* were < 0.1, indicating less phylogenetic signal than expected under BM (Table S4). 39 out of 74 of binary traits (including qualitative traits coded as binary) showed evidence of phylogenetic signal (*D* ≤ 0.5, significantly different from random; Table S3).

### Observed diversity

Taxonomic richness is lowest on the northernmost island of Hokkaido, then increases with decreasing latitude until reaching a maximum in southern Kyushu, then decreases again in smaller islands further south (Fig. 2A, Fig. S6A). Observed values of PD and FD were highly correlated with richness as expected (Fig. 2B, C), showing an initial steep slope that gradually starts to level off at higher richness (Fig. S7A, C). Observed values of RPD and RFD were not linearly related to richness; however, they showed high variance at low species richness, and low variance at high species richness (Fig. S7B, D).

**Fig 2.**
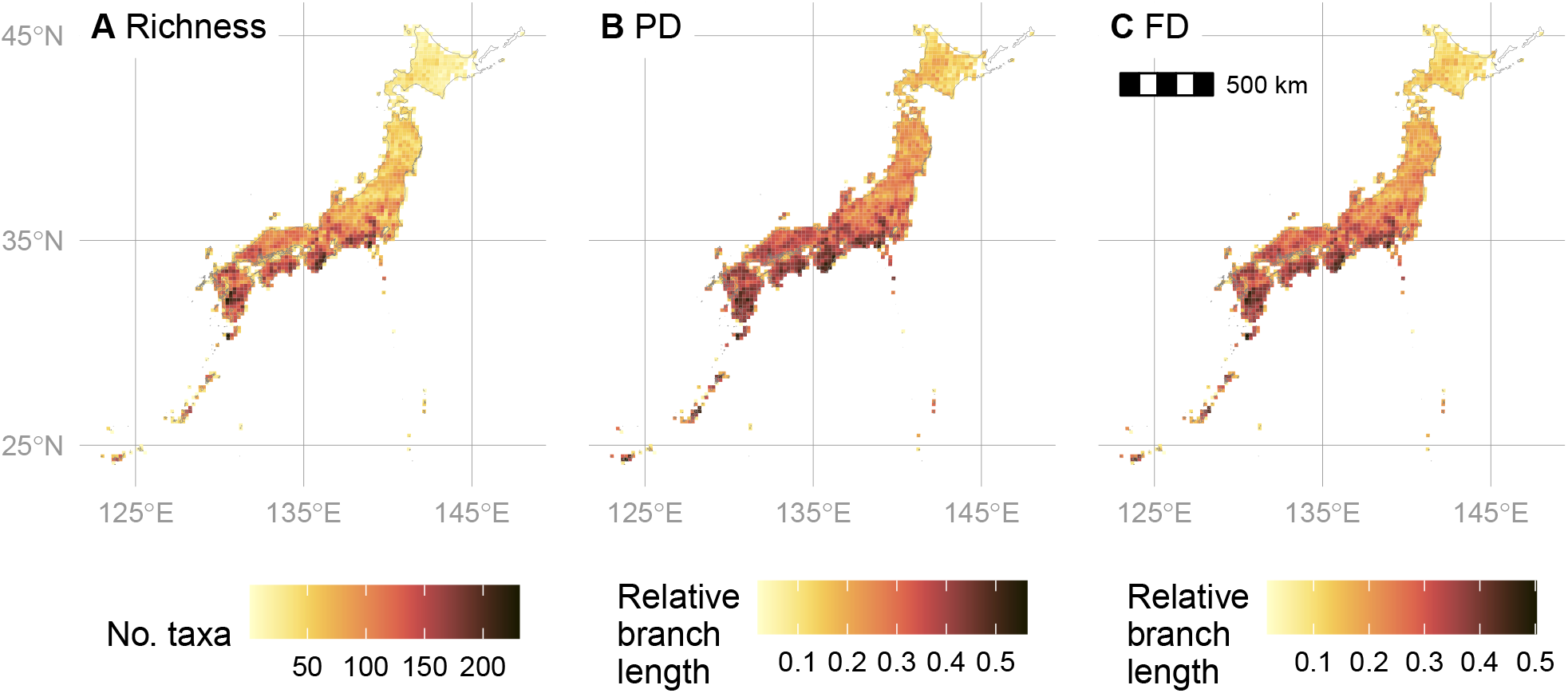
Raw (observed) biodiversity metrics of the ferns of Japan. (A) Raw taxonomic diversity (B). Raw phylogenetic diversity. (C) Raw functional diversity. For (B) and (C), raw branchlengths were transformed to relative values by dividing each branchlength by the sum of all branchlengths.

### Randomization tests for phylogenetic and functional diversity

Grid-cells with significantly low PD were almost all located on the main islands of Japan (Honshu, Kyushu, and Shikoku; Fig. 3A). A smaller number of cells had significantly high PD, and these were located mostly on small southern (Ryukyu, Izu, Ogasawara) islands or coastal areas (Kii Peninsula). Grid-cells with significantly low RPD were mostly observed in the southern main islands, particularly in and around Kyushu (Fig. 3B). Grid-cells with significantly high RPD are located both in the small southern islands and northern Honshu and Hokkaido. Results of the randomization test for functional diversity were broadly similar to phylogenetic diversity (Fig. 3C, D).

**Fig 3.**
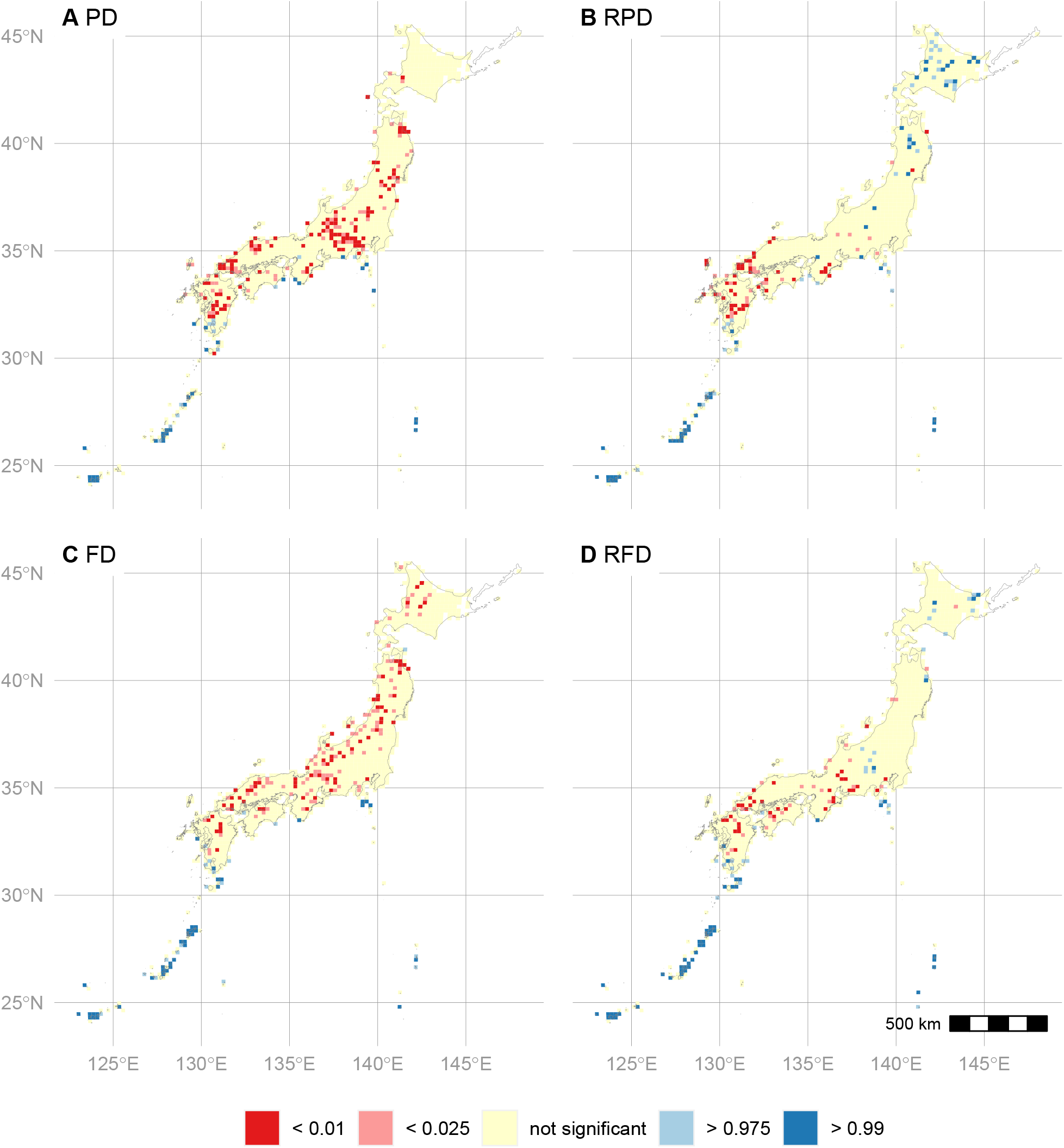
Results of randomization test for phylogenetic and functional diversity of the ferns of Japan. (A) Phylogenetic diversity (PD). (B) Relative phylogenetic diversity (RPD). (C) Functional diversity (FD). (D) Relative functional diversity (RFD). For each metric, raw values were compared to a null distribution of 999 random values, and the *P*-value calculated using a two-tailed test. For details, see Materials and Methods.

### Relationships between biodiversity and climate or reproductive mode

All models were significant relative to null models without any fixed effects (Table S5). The effect of temperature dominated (had the greatest absolute *t*-value) in environmental models (Fig. 4A–E), and % apomictic taxa dominated in reproductive models (Fig. 4F–G); temperature and % apomictic taxa were significant in each model where they were included, whereas other predictors (area, precipitation, precipitation seasonality) were not significant in some models. Richness showed a hump-shaped relationship with temperature, reaching maximum richness at intermediate temperatures (Fig. 5A). Phylogenetic and functional diversity showed either moderately (SES of PD, SES of FD) to strongly (SES of RPD, SES of RFD) hump-shaped relationships with temperature (Fig. 5B–E). SES of RPD showed a clear, linear decline with % apomictic taxa (Fig. 5G); a similar pattern was detected in SES of PD, but with wider confidence intervals (Fig. 5F). Conditional AIC was lower (indicating better model fit) in the reproductive model compared to the environmental model for SES of RPD, but not SES of PD (Table S6).

**Fig 4.**
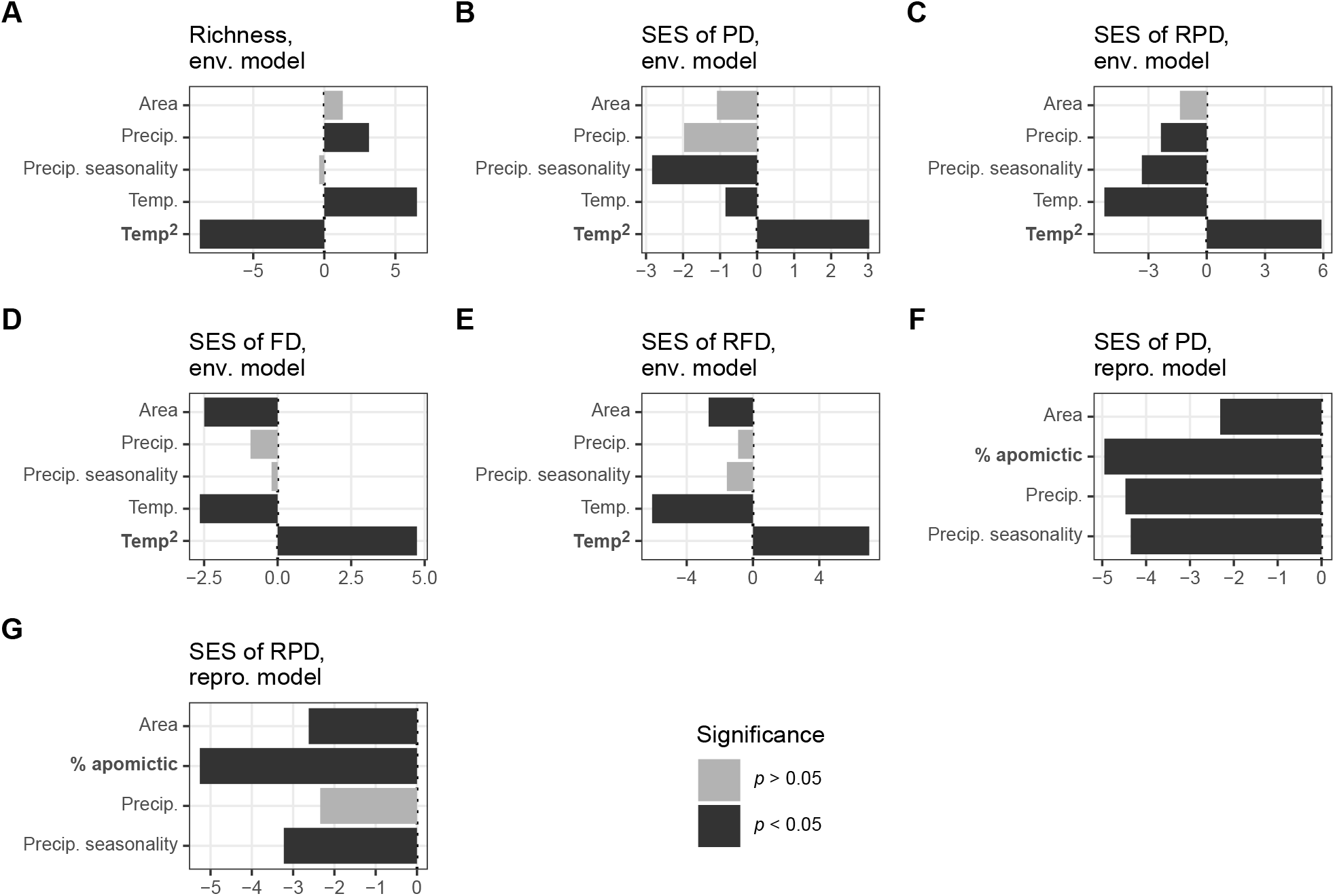
Regression coefficient *t*-values (coefficient divided by standard error) of general mixed models for the effect of environment and reproductive mode on biodiversity metrics in the ferns of Japan. (A)–(E) Environmental models (models including temperature). (F, G) Reproductive models (models including % apomictic taxa). Statistical significance assessed with a likelihood ratio test (LRT) between the full model and a model with the focal independent variable removed (null model); *P*-value indicates probability of no difference between the full and null model. Name of response variable and model type indicated above each subplot. Independent variable with greatest absolute *t*-value in bold for each model.

**Fig 5.**
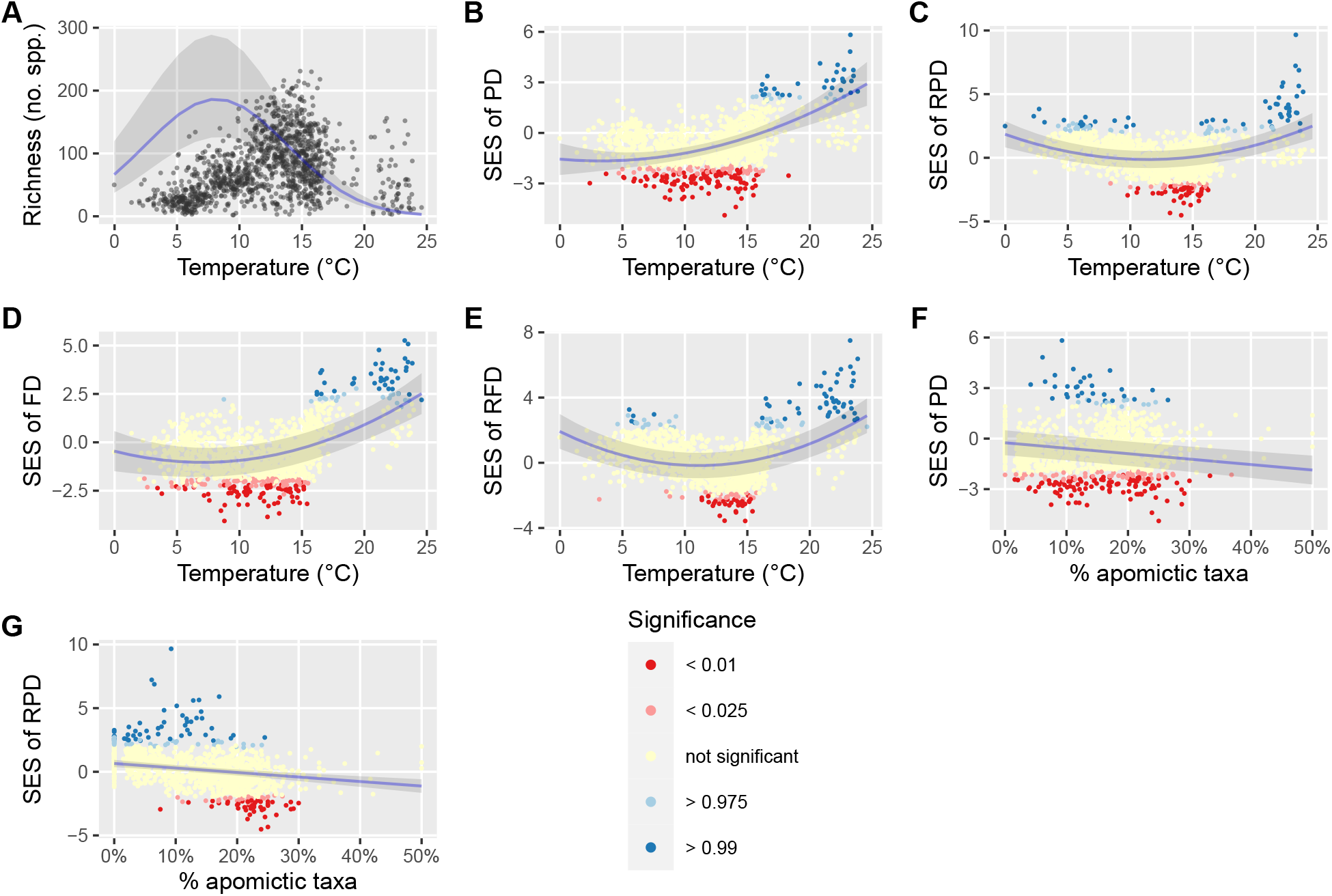
Relationship between biodiversity metrics and selected predictor variables in the ferns of Japan. (A)–(E) Environmental models (models including temperature). (F, G) Reproductive models (models including % apomictic taxa). Ribbon shows 95% confidence interval of model. Line fit with the focal predictor variable while averaging over other predictors. For phylogenetic diversity (PD), relative PD (RPD), functional diversity (FD), and relative FD (RFD), the standard effect size (SES) was calculated by comparing raw values to a null distribution of 999 random values, and significance (*P*-value) calculated using a two-tailed test (see Materials and Methods).

### Categorical analysis of neo- and paleo-endemism

Nearly all grid-cells in islands south of Kyushu (Ryukyu and Ogasawara) have significant levels of phylogenetic endemism, with low (non-significant) endemism observed elsewhere (Fig. 6). The vast majority of grid-cells with significant endemism are mixed or super-endemic. Concentrations of paleoendemism were observed in the Ryukyu archipelago, on Okinawa, Miyako, and Iriomote Islands. Small clusters of cells containing significant endemism were also observed northwest of Tokyo (Nagano prefecture), and in Hokkaido (Fig. 6). When we ran CANAPE with a reduced dataset including only taxa absolutely restricted to Japan to account for the border effect, there were fewer significant cells, but a similar pattern of significant endemism mostly occurring in the southern islands was observed (Fig. S8).

**Fig 6.**
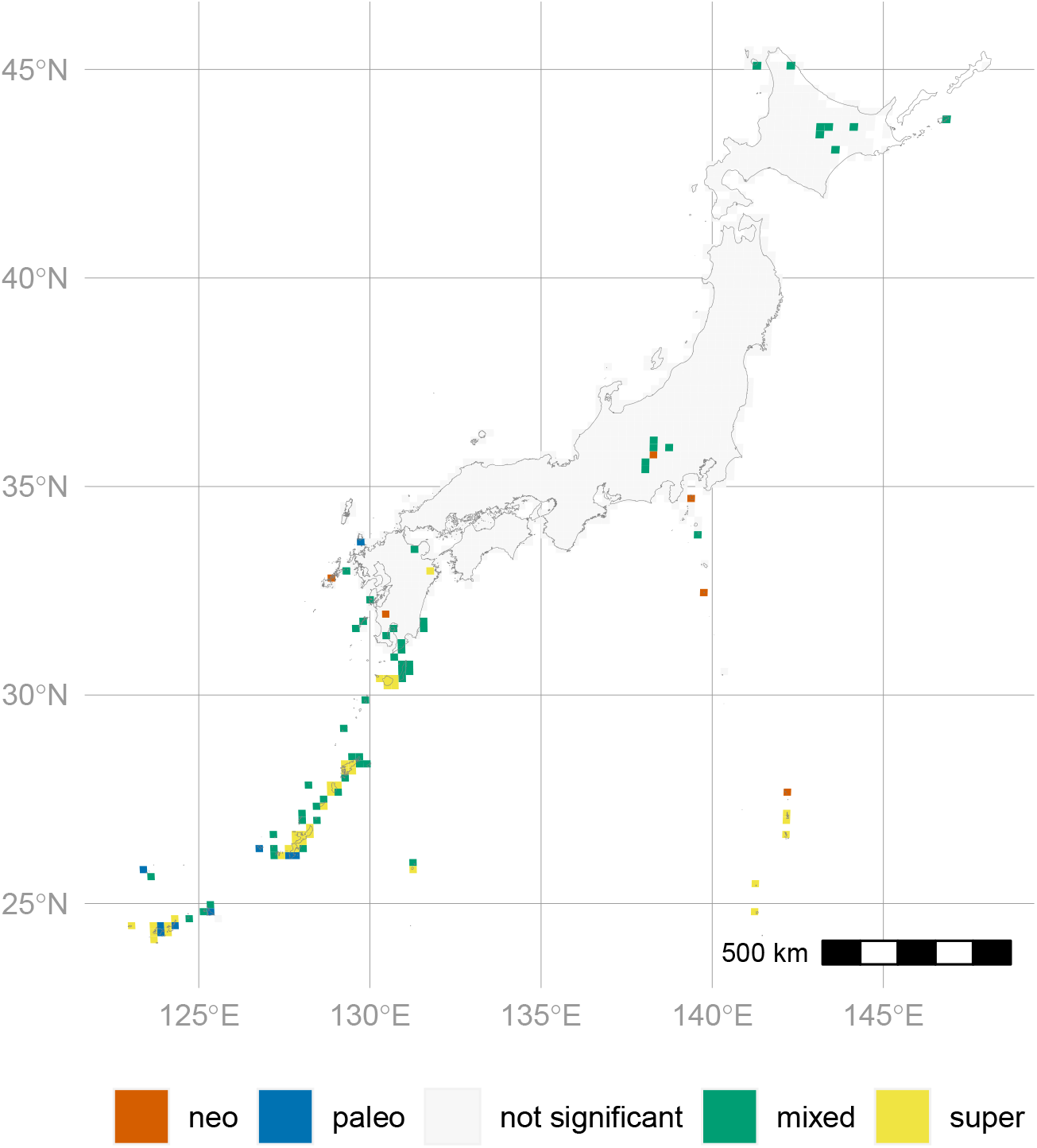
Phylogenetic endemism of the ferns of Japan measured using CANAPE (categorical analysis of neo- and paleo-endemism). ‘neo’ indicates areas with an overabundance of rare short branches; ‘paleo’ indicates areas with an overabundance of rare long branches; ‘mixed’ indicates significantly endemic sites that have neither an overabundance of short nor long branches; ‘super’ indicates highly significantly endemic sites that have neither an overabundance of short nor long branches. For details, see Materials and Methods.

### Bioregions

Clustering analysis grouped grid-cells into four or five bioregions (Fig. S9) on the basis of taxonomic or phylogenetic distances, respectively. The vast majority of grid-cells fell to a subset of these bioregions: using taxonomic distances, they include bioregion 1 (Hokkaido and high elevation areas of northern Honshu), bioregion 2 (low elevation areas of northern Honshu, southern Honshu, Kyushu, and Shikoku), bioregion 3 (Ryukyu Islands), and bioregion 4 (Ogasawara Islands) (Fig. 7A; Fig. S10A). Phylogenetic distances produced similar results, but the Ryukyu and Ogasawara Islands merged into a single bioregion (3), and bioregion 1 covered most of northern Honshu, while extending further south (Fig. 7B; Fig. S10B). The other much smaller bioregions (including only one or two grid-cells each) are likely artifacts of insufficiently low taxon richness for robust clustering and are not discussed further.

**Fig 7.**
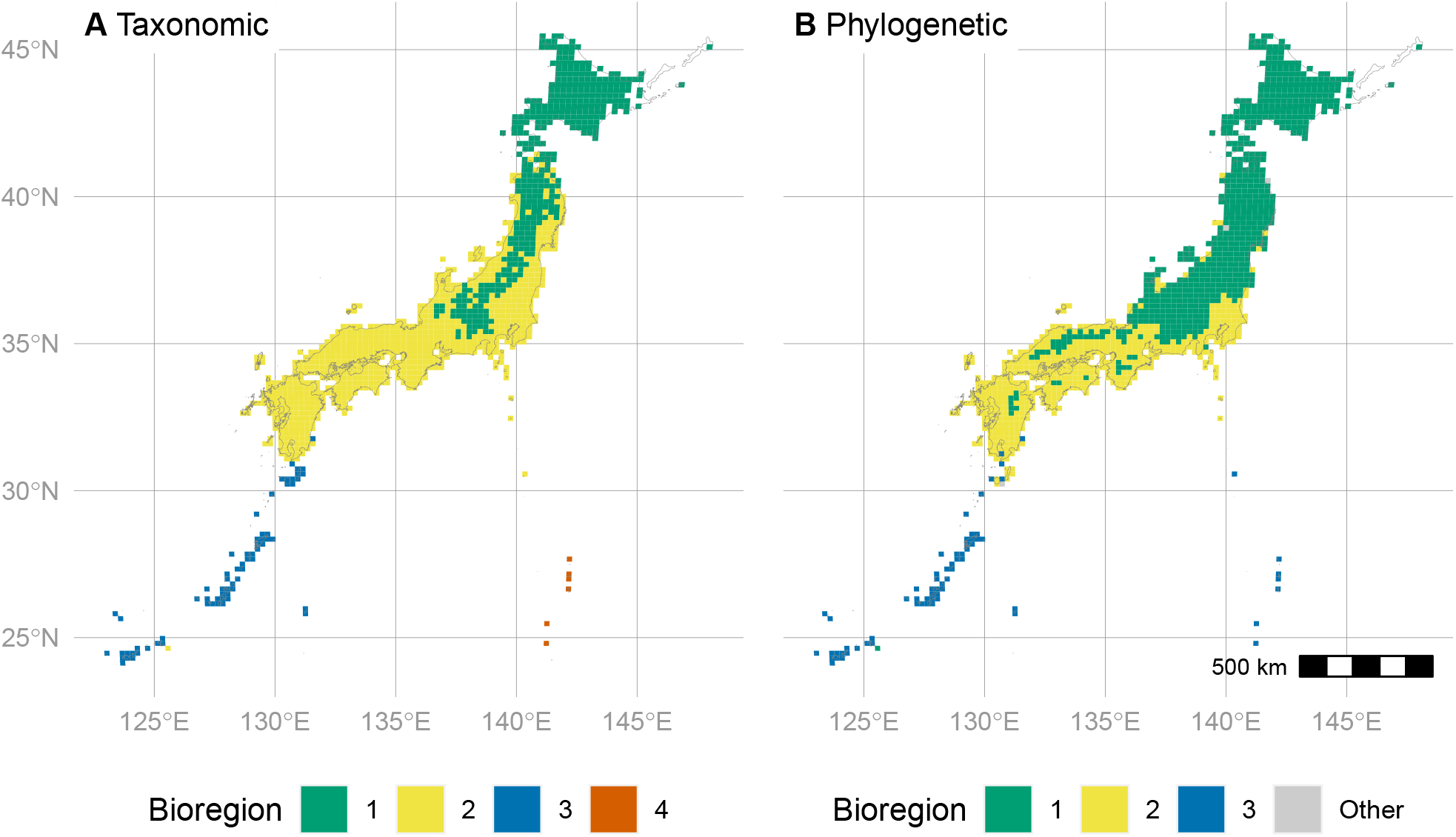
Bioregions of the ferns of Japan. (A) Taxonomic bioregions. (B) Phylogenetic bioregions. Bioregions determined by clustering taxonomic (Sørensen) or phylogenetic (PhyloSor) distances between grid-cells. Bioregions not consisting of more than two grid-cells each are lumped into the “Other” category.

Taxonomic bioregions 3 and 4 have higher mean SES values of PD, RPD, FD, and RFD than bioregions 1 and 2 (Fig. 8). Bioregions 3 and 4 also have consistently high PE *p*-scores, whereas bioregions 1 and 2 have a much wider variation (Fig. 8E). Bioregion 2 has lower RPD and RFD than bioregion 1. Comparison of biodiversity metrics across phylogenetic bioregions was similar, except that phylogenetic bioregion 3 mostly corresponds to taxonomic bioregions 3 and 4 combined.

**Fig 8.**
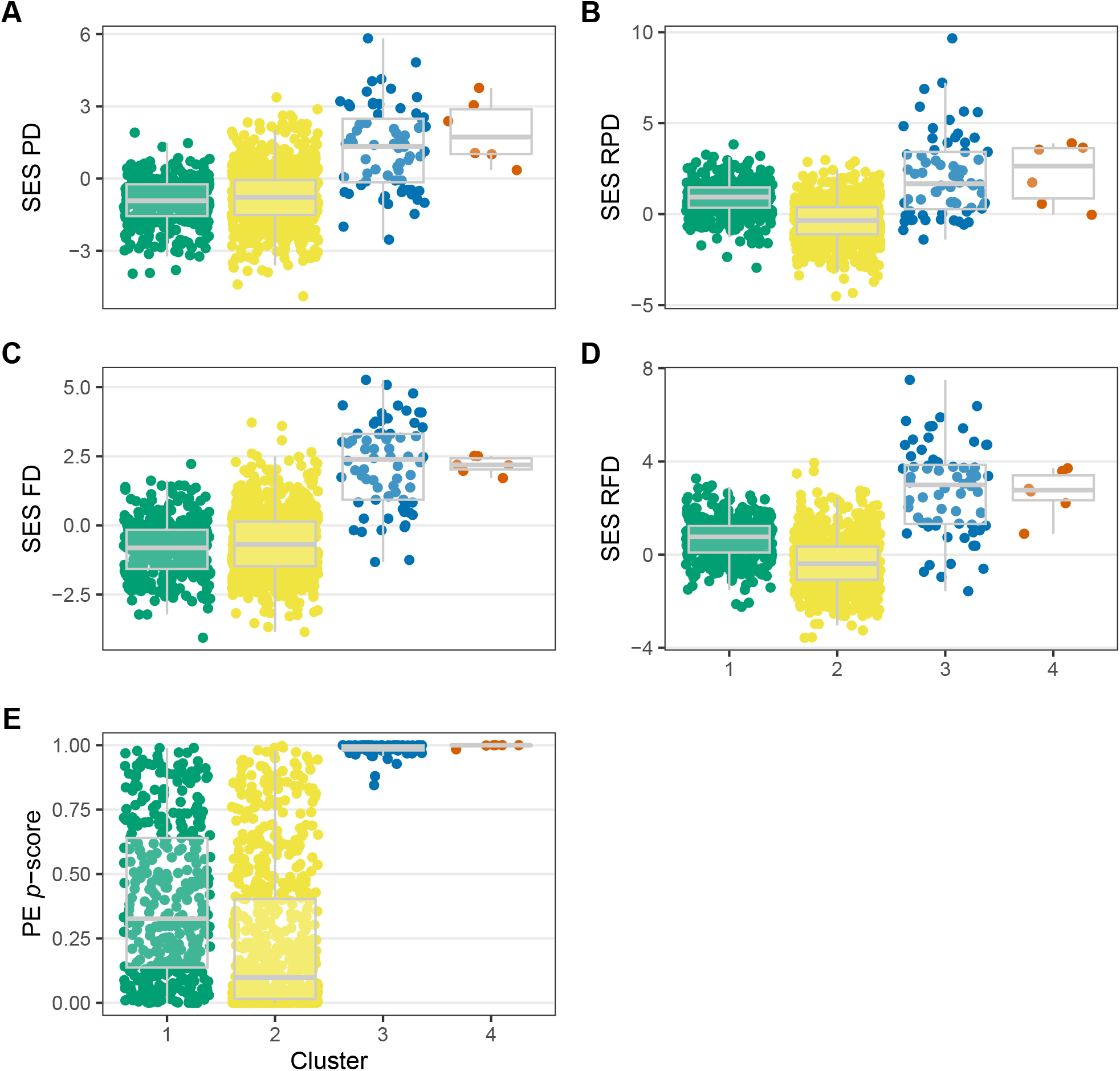
Phylogenetic and morphological diversity of the ferns of Japan by taxonomic bioregion. (A) Standard effect size (SES) of phylogenetic diversity (PD). (B) SES of relative PD (RPD). (C) SES of functional diversity (FD). (D) SES of relative FD (RFD). (E) *P*-score of observed phylogenetic endemism (PE) relative to 999 random communities. Only top four clusters with the most sites shown.

### Conservation status and threats

The percentage of protected area in grid-cells with significantly high biodiversity range from 8.6% to 23.8% total, including 2.8% to 7.0% with high status and 5.8% to 16.8% with medium status (Fig. 9A). Cells with significantly high biodiversity had a similar or greater percentage of area protected compared to the baseline rate for Japan at the medium protection status (5.9%) across all biodiversity metrics. In contrast, cells with significantly high biodiversity had a lower percentage of area protected than the baseline rate for Japan at the high protection status (3.8%) for most biodiversity metrics. Only cells with significantly high PE had a higher percentage area protected (7.0% high, 16.8% medium, 23.8% total) than the baseline rate at both levels of protection.

**Fig 9.**
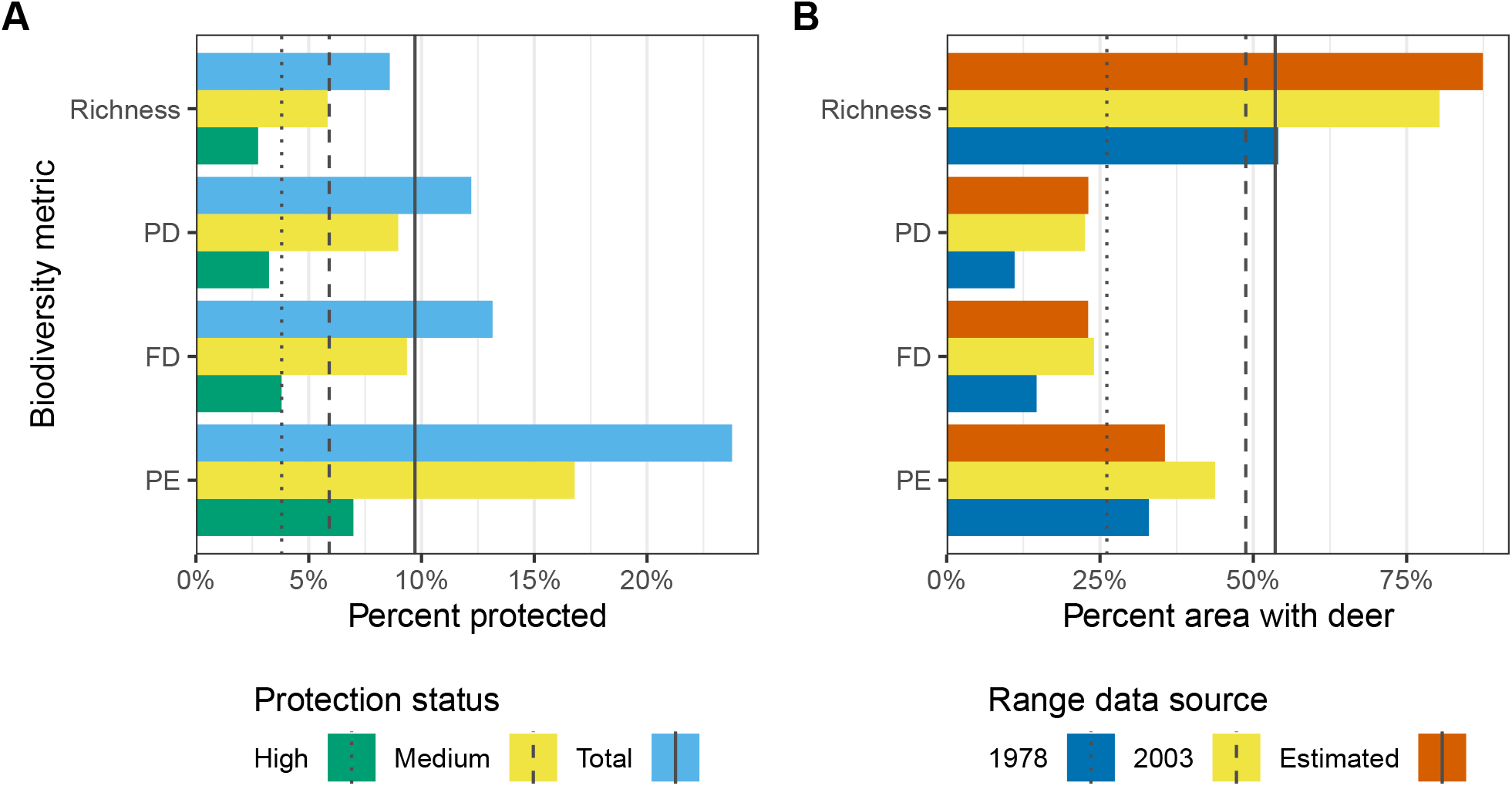
Conservation status and threat due to herbivory by deer in areas with high levels of biodiversity for ferns of Japan. (A) Percent of land area with medium or high protected status for grid-cells with significantly high biodiversity by protection status. (B) Percent of land area occupied by deer for grid-cells with significantly high biodiversity by deer range data source (1978 survey, 2003 survey, or range estimated from model based on 2003 survey data). Biodiversity metrics include taxon richness, phylogenetic diversity (PD), functional diversity (FD), and phylogenetic endemism (PE). Significance of PD, FD, and PE assessed by a one-tailed test comparing observed values to a null distribution of 999 random values; for richness, grid-cells in the top 5% considered highly diverse (see Materials and Methods). For (A), vertical lines indicate total percent area protected across Japan (baseline protection rate). For (B), vertical lines indicate total percentage of land area occupied by deer across Japan (baseline threat rate).

The range of Japanese deer in Japan increased from 98,916 km^2^ (26.1%) in 1978 to 184,671 km^2^ (48.8%) in 2003, and may be as high as 202,819 km^2^ (53.6%) according to model estimates. The percentage of area occupied by deer in grid-cells with significantly high biodiversity increased from 28.2% in 1978 to 42.7% in 2003 when averaged across biodiversity types. Cells with high FD, PD, or PE tended to have less deer occupation compared to the baseline rate (except for PE relative to 1978 levels; Fig. 9B). However, cells with high species richness consistently exceeded the baseline rate of deer presence, with up to 87.4% occupied by deer in the estimated range dataset.

## DISCUSSION

### Distribution and drivers of biodiversity

The latitudinal hump-shaped pattern of species richness observed in ferns here (Figs. 2a, S6a) is well-documented (Kusumoto et al., 2017; Ebihara and Nitta, 2019). Similar patterns have been observed in deciduous broadleaved trees and conifers (Shiono et al., 2015), and niche-modeled distributions of woody plants in Japan (Fukaya et al., 2020). Our spatial modeling analysis indicates that this is strongly linked to temperature, with maximum richness occurring at intermediate temperatures (Fig. 5A). Previous studies have identified temperature, water availability (Bickford and Laffan, 2006; Qian et al., 2012), or a combination of these (Nagalingum et al., 2015; Link-Pérez and Laffan, 2018; Qian et al., 2021) to be important drivers of spatial fern diversity. A hump-shaped pattern of richness along elevation (maximum richness at intermediate elevations) has been frequently observed in ferns on mountains in tropical or sub-tropical areas (Kessler et al., 2011; Suissa et al., 2021). Hypotheses to explain such a pattern include purely geometrical constraints as well as biological hypothesis positing maximally favorable environmental conditions at mid-elevations (Colwell et al., 2004). Another reason for the greater richness on the main islands compared to southern islands in Japan is that the main islands harbor higher elevational variation that may support a wider range of taxa than is present in the southern islands.

The overwhelming pattern characterizing phylogenetic and functional diversity (PD and FD) is the high diversity of southern subtropical islands (Ryukyu and Ogasawara Isl.) (Fig. 3A, C). This is almost certainly due to the presence of primarily tropical lineages that do not occur in other parts of the country, which lack subtropical climate; 16 genera and four families only occur south of 30.1° latitude (just south of Yakushima Isl.). The similarity in patterns of PD and FD is likely due to the moderate degree of phylogenetic signal present in the traits used in our analysis (Tables S3, S4), and suggests that at least in the ferns of Japan, phylogeny can be used as a reasonable stand-in for functional diversity. Given their high PD, it is perhaps not surprising that southern islands also host high amounts of phylogenetic endemism (PE); indeed, the vast majority of cells with significant levels of PE, as well as paleoendemic cells, are located in the southern islands (Figs. 6, S8). This is clearly due to the small land area of these islands: small area combined with high PD is expected to lead to high PE.

Another striking pattern revealed here is the predominance of long branches in the northern half of Honshu and the southern islands, and the predominance of short branches in southern Honshu, Shikoku, and Kyushu (Fig. 3B). The long branches in the southern islands are likely being driven by the tropical lineages restricted to this area; those in the north may be due to refugial lineages that are distantly related to others and become more species rich at high latitudes, such as *Equisetum* (Fig. S11A). The preponderance of short branches in southern Honshu, Shikoku, and Kyushu may be due to the radiation of certain species-rich genera there such as *Deparia* (Fig. S11B) and *Dryopteris* (Fig. S11D).

Furthermore, we identified % apomictic taxa as a strong driver of SES of RPD (Figs. 4G, 5G), and to a slightly lesser extent SES of PD (Figs. 4F, 5F). Indeed, *Dryopteris* is one such genus with a high rate of apomictic taxa (Fig. S12), and other genera with high rates of apomixis (*Cyrtomium, Pteris*; Fig. S12) also reach greatest richness in the southern main islands (Fig. S11). This is consistent with the hypothesis that communities with high proportions of asexual taxa have lower PD, and is the first time to our knowledge that this has been demonstrated in ferns. We verified that this relationship is also observed when analyzed with a phylogeny with branch lengths in units of genetic change (*ie*, the output of the maximum likelihood phylogenetic analysis, before dating), a more direct measure of genetic diversity (Fig. S13). While temperature is also correlated with % apomictic taxa and therefore cannot be discounted, conditional AIC indicates a better fit of the reproductive model than the environmental model for SES of RPD (Table S6). Ultimately, it is likely that temperature, in combination with other environmental conditions, is driving the predominance of apomictic taxa: Tanaka et al. (2014) also found that the proportion of apomictic taxa reaches a maximum at ca. 34.8 °N latitude and increases with temperature in Japan (their study was limited to the main islands and did not include the smaller southern islands). Tanaka et al. (2014) asserted that one limiting factor affecting establishment of apomictic ferns at high latitudes or cold climates is the large cell size of apomicts, which are typically polyploid. As fern gametophytes are only a single cell-layer in thickness with no cuticle or stomata, it is expected they would be particularly susceptible to freezing damage induced by larger cells with higher water content; however, no experimental studies have tested these hypotheses to our knowledge.

Water limitation is thought to play an important role in promoting apomixis in ferns especially in deserts (Grusz et al., 2021), since ferns depend on water for transfer of sperm to egg, and apomictic plants would be able to reproduce in the absence of water. A similar argument has been made to assert that areas with seasonal monsoons are linked to increased rates of apomixis in ferns (Liu et al., 2012; Tanaka et al., 2014; Picard et al., 2021), but in Japan, although the areas including high percentages of apomictic taxa are monsoonal, the degree of seasonal water limitation does not approach that of a desert. The monsoon hypothesis was not supported by our results, which did not show a particularly strong effect of precipitation seasonality in most models (Fig. 4), nor a correlation with % apomictic taxa (Table S2).

The variation in distribution of different biodiversity metrics is reflected by the bioregions analysis, which tended to group cells with similar biodiversity values (Fig. 8). Interestingly, the bioregions identified here correspond roughly to major vegetation zones previously categorized on an *ad-hoc* basis; bioregion 1 corresponds to the cool or subartic deciduous type, bioregion 2 to the warm evergreen type, and bioregions 3 and 4 to the subtropical rainy type (Shimizu, 2014). Furthermore, the border between bioregions 1 and 2 approximately corresponds to the “*Gleichenia japonica* Line” proposed by Nakaike (1983), which he proposed based on the distribution of *Diplopterygium glaucum* (Houtt.) Nakai (= *Gleichenia japonica* Spreng.) as an indicator of evergreen broad-leaf forests in Japan. This suggests that ferns are useful as bioindicators in Japan. Although some studies have emphasized the importance of deep-sea straits (the Tsugaru Strait separating Hokkaido from Honshu and the Tokara Strait separating the southern islands from Kyushu) in structuring biological communities in Japan (Millien-Parra and Jaeger, 1999; Kubota et al., 2014), these did not play a major role in structuring fern bioregions. The Tsugaru Strait had no relation to any of the bioregions, and although the Tokara Strait somewhat splits phylogenetic bioregions 2 and 3, there is some overlap between them on Yakushima and Amami Isl. (Fig. 7B). The weak effect of these straits as barriers makes sense given the presumably high dispersal abilities of ferns, which disperse via tiny spores carried on the wind (Tryon, 1970). Rather, fern bioregions in Japan broadly separate along gradients of temperature and precipitation; in particular, precipitation splits taxonomic bioregion 3 (mostly the Ryukyu islands; wetter) from bioregion 4 (mostly the Ogasawara islands; drier) (Fig. S14).

### Conservation status and threats

We found that grid-cells with significantly high amounts of biodiversity are similarly or better protected compared to the baseline protection rate at the medium protection level across all biodiversity metrics, and better than the baseline rate at the high protection level for PE (Fig. 9A). The high protection status of grid-cells with high PE may be due to an emphasis on conserving endemic plant and animal species in Japanese southern islands such as Iriomote and Amami (Japan Ministry of the Environment, 2021a), which include many grid-cells with high PE (Fig. S15D). One possible reason for the generally high level of protection overall is that in the past, some ferns have been included in lists of rare and endemic plants for consideration in designation of conservation areas (A. Ebihara, pers. comm.; Japan Ministry of the Environment, 2021a). Also, (as suggested previously) ferns may serve as bioindicators reflecting overall levels of biodiversity, particularly for other plant groups.

Despite the good news that ferns enjoy relatively strong protection status in Japan, we must bear in mind that there are other threats posed that conservation zones cannot ameliorate, the most obvious being climate change. One other such threat is herbivory by native Japanese deer, *Cervus nippon* (Minamitani, 2005; Yahara, 2006; Hattori et al., 2010; Japan Ministry of the Environment, 2022). The rapid increase in population size observed in surveys from 1978 to 2003 is likely due to a combination of climate change and a decrease in human hunters and natural predators (Takatsuki, 2009). At dense populations, the deer can prevent forest regeneration and result in plant communities dominated by a limited number of species unpalatable to deer (Takatsuki, 2009), although lower population sizes may promote plant diversity as per the intermediate disturbance hypothesis (Nishizawa et al., 2016). They have caused extensive damage even within national parks (Tokita, 2006), which have the highest protection status. Although various efforts to prevent deer herbivory have been deployed such as culling and exclusion fences (Takatsuki, 2009), much work remains to be done to save vulnerable plant populations.

Our analysis shows that the threat to fern biodiversity posed by Japanese deer differs strongly by biodiversity type; although grid-cells with significantly high FD, PD, and PE tend to have lower rates of deer occupancy compared to the baseline rate in recent surveys or modeled data, grid-cells with high richness experience much greater co-occurrence with deer than the baseline rate (Fig. 9B). This pattern is due to the fact that Japanese deer do not occur in the southern islands, which house high amounts of PD and PE; rather, areas with high species richness are primarily located in southern Honshu, Shikoku, and Kyushu, where deer expansion has been intense (Fig. S15A). This result highlights the importance of using multiple metrics of biodiversity for establishing conservation priorities.

## CONCLUSIONS

Our study integrates diverse, thoroughly collected datasets to characterize the spatial biodiversity of Japanese ferns and gain insight into the processes generating these patterns. We conclude by summarizing some caveats that must be kept in mind when interpreting our results, while also suggesting avenues for future research.

First, the data include specimens collected over a wide time period, so they do not necessarily represent the current distribution of ferns in Japan. Furthermore, the collections are not evenly collected throughout the country, but rather some areas have many more specimens than others (Fig. S5). Indeed, one promising avenue of research may be to take advantage of the most densely sampled areas to investigate changes in abundance over time.

The extreme evolutionary distinctiveness of the southern islands is both a major feature of the Japanese fern flora and a challenge for analysis. The tropical lineages occurring only in these islands are likely responsible for the comparatively low (*ie*, clustered) values of PD observed on the main islands (Fig. 3A). One way to account for this bimodal distribution pattern would be to use a restricted null model that does not allow all taxa to disperse anywhere within the study area (Mishler et al., 2020). However, ferns are known to have high dispersal abilities (Tryon, 1970; Barrington, 1993; Moran and Smith, 2001). In our dataset, the taxon with the greatest latitudinal distribution (*Pteridium aquilinum* (L.) Kuhn subsp. *japonicum* (Nakai) A. et S. Löve) spans nearly the entire region (24.3°N to 45.46°N), and 16 taxa have latitudinal distributions spanning the Tokara strait in the south (29°N) to the Tsugaru strait in the north (41.5°N). Therefore, it seems reasonable to assume that ferns are capable of dispersing across the entire study area.

Furthermore, it should be noted that the southern islands are all low-altitude (not exceeding 1,000 m). If high elevation habitats were available, the southern islands would be expected to share more species in common with areas further north. For example, Taiwan, which is just east of the southernmost Japanese islands and harbors high mountains (> 3,000 m), shares many fern species with the main Japanese islands that are not found in southern Japanese islands. This supports that the sharp taxonomic turnover between southern and main islands is more likely due to ecological factors than dispersal limitation.

One element of biodiversity that we did not consider here is hybrids. The Japanese fern flora includes 374 nothotaxa (Ebihara and Nitta, 2019). Although hybrids are often thought of as evolutionary “dead-ends”, in some cases they are able to interbreed with diploids and thus contribute to diversification (Barrington et al., 1989). Future studies should consider the distribution of hybrids in Japan and their possible effects on biodiversity. Methods for calculating PD would need to be modified to take into account multiple lineages contributing to one terminal taxon.

Finally, a long under-appreciated aspect of fern ecology is the role of the gametophyte. Unlike seed plants, ferns have gametophytes that are capable of growing independently from the sporophyte. Futhermore, the two stages of the life cycle have vastly different morphology and ecophysiology, and may even occur over partially to completely disjunct ranges (reviewed in Pinson et al., 2017). Here, as in the vast majority of fern ecology studies, we consider only the sporophyte stage. However, recent studies in Japan (Ebihara et al., 2013) and elsewhere (Nitta et al., 2017) have revealed different patterns in the community structure of gametophytes and sporophytes. The comprehensive molecular sampling of sporophytes in Japan makes this area ideal for conducting high-throughput DNA sequencing analyses to compare patterns of biodiversity between life stages in ferns across large spatial scales.

## ACKNOWLEDGEMENTS

The authors thank Members of the Iwasaki Lab at the University of Tokyo and two anonymous reviewers for providing comments that improved the manuscript. Shawn Laffan provided advice about CANAPE. This study was supported by JSPS KAKENHI Grant Number 16H06279. The authors are grateful to members of the Nippon Fernist Club for their cooperation and efforts towards quantifying and conserving the biodiversity of the ferns of Japan.

## AUTHOR CONTRIBUTIONS

J.H.N., B.M., and A.E. conceived the ideas; A.E. provided the data; J.H.N. analyzed the data; W.I provided resources; J.H.N. wrote the manuscript with input from all co-authors.

## DATA AVAILABILITY STATEMENT

Code to replicate all analyses, figures, and this manuscript are available at https://github.com/joelnitta/japan_ferns_spatial_phy. A Docker image to run the code is available at https://hub.docker.com/r/joelnitta/japan_ferns_spatial_phy. Data files needed to run the analysis and selected results files are available from Figshare at http://doi.org/10.6084/m9.figshare.16655263.

## SUPPORTING INFORMATION

Additional supporting information may be found online in the Supporting Information section at the end of the article.

**Appendix S1**. Supplementary tables and figures, including Tables S1–S6 and Figs. S1–S16.

**Table S1**. Spatial autocorrelation in biodiversity metrics (richness, SES of PD, SES of RPD, SES of FD, SES of RFD) and independent variables (environmental variables and % apomictic taxa) as measured with Moran’s *I*.

**Table S2**. Results of modified *t*-test for correlation between variables while taking into account spatial position.

**Table S3**. Phylogenetic signal in quantitative (binary) functional traits of the ferns of Japan.

**Table S4**. Phylogenetic signal in continuous functional traits of the ferns of Japan.

**Table S5**. Likelihood ratio test (LRT) between full model and null model (model only including the spatial Matérn correlation matrix).

**Table S6**. Model fit as measured with conditional Akaike Information Criterion (cAIC).

**Fig. S1**. Species collection curve for the ferns of Japan.

**Fig. S2**. Effect of grain size on sampling redundancy.

**Fig. S3**. Reproductive mode and environmental data in 20 km grid cells across Japan.

**Fig. S4**. Time tree of native Japanese ferns and lycophytes, excluding nothotaxa.

**Fig. S5**. Observed number of specimens and sampling redundancy per 20 km grid cell in the ferns of Japan.

**Fig. S6**. Raw taxonomic diversity, phylogenetic diversity, and functional diversity of the ferns of Japan plotted by latitude.

**Fig. S7**. Relationships between observed functional and phylogenetic diversity and taxonomic richness in the ferns of Japan.

**Fig. S8**. Phylogenetic endemism of the ferns of Japan measured using CANAPE (categorical analysis of neo- and paleo-endemism), restricted dataset including only taxa endemic to Japan.

**Fig. S9**. Selection of *k* for bioregions.

**Fig. S10**. Bioregion dendrograms.

**Fig. S11**. Map of species richness for selected genera of ferns of Japan.

**Fig. S12**. Rate of apomixis in genera of Japanese ferns with > 1 apomictic taxon.

**Fig. S13**. Relationship between phylogenetic diversity (PD) and relative PD and % apomictic taxa, models inferred using phylogeny with branchlengths in units of expected genetic change (untransformed ML tree).

**Fig. S14**. Scatterplots of grid cell membership in taxonomic or phylogenetic bioregions arranged by mean annual temperature and annual precipitation.

**Fig. S15**. Maps showing overlap of grid-cells with significantly high biodiversity for the ferns of Japan and protected areas.

**Fig. S16**. Maps showing overlap of grid-cells with significantly high biodiversity for the ferns of Japan and distribution of Japanese deer (*Cervus nippon*).

**Appendix S2**. Exploratory data analysis for spatial modeling.

