## Appendix S1 for "Spatial phylogenetics of Japanese ferns: Patterns, processes, and implications for conservation"

### Appendix S1: Supporting Tables and Figures

| Variable | *I* | *P* |
|---|---|---|
| Area | 0.354 | 0.001 |
| % apomictic taxa | 0.376 | 0.001 |
| Precipitation | 0.304 | 0.001 |
| Precipitation seasonality | 0.355 | 0.001 |
| Temperature | 0.344 | 0.001 |
| Richness | 0.265 | 0.001 |
| SES of PD | 0.079 | 0.001 |
| SES of RPD | 0.162 | 0.001 |
| SES of FD | 0.095 | 0.001 |
| SES of RFD | 0.127 | 0.001 |

**Table S1**. Spatial autocorrelation in biodiversity metrics (richness, SES of PD, SES of RPD, SES of FD, SES of RFD) and independent variables (environmental variables and % apomictic taxa) as measured with Moran's *I*. Significant *I* > 0 indicates autocorrelation.

| Variable 1 | Variable 2 | *P* | *r* | *f* | df |
|---|---|---|---|---|---|
| % apomictic taxa | Precipitation | 0.0956 | 0.494 | 0.3224 | 10.4 |
| % apomictic taxa | Precipitation seasonality | 0.1561 | 0.491 | 0.3172 | 7.8 |
| **% apomictic taxa** | **Temperature** | **0.0160** | **0.648** | **0.7235** | **11.1** |
| **% apomictic taxa** | **Temperature seasonality** | **0.0195** | **-0.482** | **0.3034** | **21.1** |
| Area | % apomictic taxa | 0.3998 | 0.229 | 0.0554 | 13.6 |
| Area | Precipitation | 0.6802 | -0.068 | 0.0046 | 37.6 |
| Area | Precipitation seasonality | 0.5409 | 0.178 | 0.0327 | 12.1 |
| Area | Temperature | 0.4594 | -0.068 | 0.0047 | 117.6 |
| Area | Temperature seasonality | NaN | 0.229 | 0.0555 | -128.4 |
| Precipitation | Precipitation seasonality | 0.2645 | 0.285 | 0.0885 | 15.2 |
| Precipitation | Temperature | 0.0704 | 0.582 | 0.5115 | 8.4 |
| **Precipitation** | **Temperature seasonality** | **0.0413** | **-0.571** | **0.4833** | **11.0** |
| Precipitation seasonality | Temperature | 0.1674 | 0.281 | 0.0859 | 23.6 |
| **Precipitation seasonality** | **Temperature seasonality** | **0.0333** | **-0.216** | **0.0490** | **95.3** |
| **Temperature** | **Temperature seasonality** | **0.0035** | **-0.898** | **4.1778** | **5.6** |

**Table S2**. Results of modified *t*-test for correlation between variables while taking into account spatial position. Significantly correlated variables at $P < 0.05$ in bold.

| Trait | *D* | *N* present | *N* absent | *P*(random) | *P*(Brownian) |
|---|---|---|---|---|---|
| indusium_present | -0.207 | 491 | 172 | 0.000 | 0.980 |
| margin_acutely_serrate | 0.075 | 32 | 631 | 0.000 | 0.347 |
| margin_biserrate | -0.133 | 7 | 656 | 0.000 | 0.615 |
| margin_crenate | 0.449 | 93 | 570 | 0.000 | 0.000 |
| margin_dentate | 0.051 | 52 | 611 | 0.000 | 0.372 |
| margin_entire | 0.166 | 317 | 346 | 0.000 | 0.015 |
| margin_minutely_serrate | 0.399 | 29 | 634 | 0.000 | 0.017 |
| margin_notched | 0.416 | 15 | 648 | 0.000 | 0.081 |
| margin_serrate | 0.235 | 198 | 465 | 0.000 | 0.002 |
| margin_undulate | 0.382 | 155 | 508 | 0.000 | 0.000 |
| shape_broadlly_pentangular | -0.151 | 17 | 646 | 0.000 | 0.709 |
| shape_broadly_lanceolate | 0.923 | 19 | 644 | 0.117 | 0.000 |
| shape_broadly_oblong_lanceolate | 0.624 | 36 | 627 | 0.000 | 0.000 |
| shape_broadly_ovate | 0.815 | 11 | 652 | 0.033 | 0.001 |
| shape_broadly_pentagonal_triangular | 0.371 | 6 | 657 | 0.000 | 0.232 |
| shape_broadly_triangular | 0.347 | 24 | 639 | 0.000 | 0.051 |
| shape_broadly_triangular_lanceolate | 0.776 | 19 | 644 | 0.001 | 0.000 |
| shape_broadly_triangular_oblong | 0.965 | 2 | 661 | 0.425 | 0.171 |
| shape_broadly_triangular_ovate | 0.640 | 35 | 628 | 0.000 | 0.000 |
| shape_broady_ovate_lanceolate | 0.457 | 34 | 629 | 0.000 | 0.006 |
| shape_elliptical | 0.276 | 13 | 650 | 0.000 | 0.175 |
| shape_fan_shape | 0.424 | 6 | 657 | 0.001 | 0.174 |
| shape_lanceolate | 0.769 | 66 | 597 | 0.000 | 0.000 |
| shape_linear | 0.212 | 12 | 651 | 0.000 | 0.264 |
| shape_linear_lanceolate | 0.577 | 12 | 651 | 0.000 | 0.026 |
| shape_linear_oblanceolate | 0.249 | 6 | 657 | 0.000 | 0.303 |
| shape_narrowly_elliptical | 0.626 | 9 | 654 | 0.000 | 0.037 |
| shape_narrowly_lanceolate | 0.652 | 32 | 631 | 0.000 | 0.000 |
| shape_narrowly_oblanceolate | 0.820 | 5 | 658 | 0.074 | 0.039 |
| shape_narrowly_oblong | 0.558 | 26 | 637 | 0.000 | 0.003 |
| shape_narrowly_oblong_lanceolate | 0.850 | 14 | 649 | 0.029 | 0.000 |
| shape_narrowly_ovate | 0.686 | 12 | 651 | 0.002 | 0.011 |
| shape_narrowly_ovate_elliptical | 0.133 | 6 | 657 | 0.000 | 0.410 |
| shape_narrowly_ovate_lanceolate | 1.092 | 5 | 658 | 0.756 | 0.000 |
| shape_narrowly_ovate_oblong | 0.918 | 7 | 656 | 0.182 | 0.005 |
| shape_narrowly_triangular_oblong | 0.996 | 6 | 657 | 0.438 | 0.005 |
| shape_narrowly_triangular_ovate | 0.997 | 7 | 656 | 0.449 | 0.001 |
| shape_oblanceolate | 0.929 | 5 | 658 | 0.276 | 0.017 |
| shape_oblong | 0.730 | 9 | 654 | 0.014 | 0.009 |
| shape_oblong_lanceolate | 0.629 | 67 | 596 | 0.000 | 0.000 |
| shape_obovate_elliptical | -0.565 | 3 | 660 | 0.000 | 0.783 |
| shape_obovate_oblong | -0.526 | 3 | 660 | 0.000 | 0.776 |

*(continued)*

| Trait | *D* | *N* present | *N* absent | *P*(random) | *P*(Brownian) |
|---|---|---|---|---|---|
| shape_ovate | 0.830 | 45 | 618 | 0.005 | 0.000 |
| shape_ovate_elliptical | 0.648 | 14 | 649 | 0.000 | 0.006 |
| shape_ovate_lanceolate | 0.624 | 23 | 640 | 0.000 | 0.001 |
| shape_ovate_oblong | 0.354 | 26 | 637 | 0.000 | 0.042 |
| shape_pentagonal_ovate | 1.022 | 4 | 659 | 0.534 | 0.015 |
| shape_pentangular | 1.091 | 4 | 659 | 0.725 | 0.010 |
| shape_triangular | 0.488 | 35 | 628 | 0.000 | 0.002 |
| shape_triangular_elliptical | -1.306 | 3 | 660 | 0.000 | 0.926 |
| shape_triangular_lanceolate | 0.807 | 19 | 644 | 0.007 | 0.000 |
| shape_triangular_oblong | 0.864 | 26 | 637 | 0.016 | 0.000 |
| shape_triangular_ovate | 0.654 | 115 | 548 | 0.000 | 0.000 |
| texture_coriaceous | 0.376 | 47 | 615 | 0.000 | 0.005 |
| texture_extremely_hard_papyraceous | -0.270 | 3 | 659 | 0.001 | 0.619 |
| texture_filmy | -0.533 | 40 | 622 | 0.000 | 1.000 |
| texture_hard_herbaceous | 0.556 | 16 | 646 | 0.000 | 0.017 |
| texture_hard_papyraceous | 0.442 | 56 | 606 | 0.000 | 0.002 |
| texture_hardish_herbaceous | 0.375 | 19 | 643 | 0.000 | 0.063 |
| texture_hardish_papyraceous | 0.068 | 26 | 636 | 0.000 | 0.379 |
| texture_herbaceous | 0.393 | 160 | 502 | 0.000 | 0.000 |
| texture_papyraceous | 0.392 | 94 | 568 | 0.000 | 0.000 |
| texture_soft_coriaceous | 0.464 | 30 | 632 | 0.000 | 0.012 |
| texture_soft_papyraceous | -0.174 | 8 | 654 | 0.000 | 0.680 |
| texture_supple_coriaceous | -0.174 | 9 | 653 | 0.000 | 0.677 |
| texture_thick_coriaceous | 0.944 | 3 | 659 | 0.385 | 0.075 |
| texture_thick_fleshy | 1.340 | 3 | 659 | 0.923 | 0.012 |
| texture_thick_herbaceous | 0.562 | 34 | 628 | 0.000 | 0.000 |
| texture_thick_papyraceous | 0.296 | 25 | 637 | 0.000 | 0.083 |
| texture_thickish_herbaceous | 0.739 | 11 | 651 | 0.006 | 0.004 |
| texture_thickish_papyraceous | -0.217 | 3 | 659 | 0.001 | 0.616 |
| texture_thin_coriaceous | 0.684 | 21 | 641 | 0.000 | 0.000 |
| texture_thin_herbaceous | 0.392 | 39 | 623 | 0.000 | 0.009 |
| texture_thin_papyraceous | 0.460 | 15 | 647 | 0.000 | 0.046 |

**Table S3**. Phylogenetic signal in quantitative (binary) functional traits of the ferns of Japan. *D*, Fritz and Purvis' *D*; *N* (present), number of times a trait was observed present; *N* (absent), number of times a trait was observed absent; *P*(random), probability of obtaining observed trait distribution given random distribution of traits; *P*(Brownian), probability of obtaining observed trait distribution given random evolution by Brownian Motion. Presence or absence of indusium was originally a binary trait; other traits (margin, shape, texture) are qualitative traits scored in binary format. Values of *D* rounded to three decimal places.

| Trait | *K* | *P*(*K*) | λ | *P*(λ) | *N* |
|---|---|---|---|---|---|
| frond_width | 0.038 | 0.001 | 0.97 | 1.6e-190 | 646 |
| stipe_length | 0.035 | 0.001 | 1.00 | 1.0e-147 | 596 |
| number_pinna_pairs | 0.012 | 0.001 | 0.98 | 3.1e-81 | 556 |

**Table S4**. Phylogenetic signal in continuous functional traits of the ferns of Japan. *K*, Blomberg's *K*; λ, Pagel's λ.

| Response variable | $\chi^2$ | df | LogL(null) | LogL(full) | *P* |
|---|---|---|---|---|---|
| **Environmental** | | | | | |
| Richness | 218 | 5 | -5867 | -5759 | 0.0e+00 |
| SES of FD | 63 | 5 | -1704 | -1672 | 2.5e-12 |
| SES of PD | 81 | 5 | -1749 | -1708 | 5.6e-16 |
| SES of RFD | 58 | 5 | -1748 | -1719 | 2.6e-11 |
| SES of RPD | 51 | 5 | -1800 | -1775 | 8.4e-10 |
| **Reproductive** | | | | | |
| SES of PD | 63 | 4 | -1749 | -1717 | 5.4e-13 |
| SES of RPD | 48 | 4 | -1800 | -1776 | 7.5e-10 |

**Table S5**. Likelihood ratio test (LRT) between full model and null model (model only including the spatial Matérn correlation matrix). Higher (less negative) log likelihood indicates better fit. Full environmental models included the effects of temperature, precipitation, precipitation seasonality, and area. Full reproductive models included the effects of % apomictic taxa, precipitation, precipitation seasonality, and area. PD, phylogenetic diversity; RPD, relative phylogenetic diversity; FD, functional diversity; RFD, relative functional diversity; SES, standard effect size.

| Response variable | cAIC | df | LogL |
|---|---|---|---|
| **Environmental** | | | |
| SES of PD | -6208 | 0.208 | -1708 |
| SES of RPD | 1854 | 142.654 | -1775 |
| **Reproductive** | | | |
| SES of PD | 2197 | 229.593 | -1717 |
| SES of RPD | -8598 | 0.027 | -1776 |

**Table S6**. Model fit as measured with conditional Akaike Information Criterion (cAIC). Environmental models included the effects of temperature, precipitation, precipitation seasonality, and area. Reproductive models included the effects of % apomictic taxa, precipitation, precipitation seasonality, and area. Lower cAIC indicates better fit.

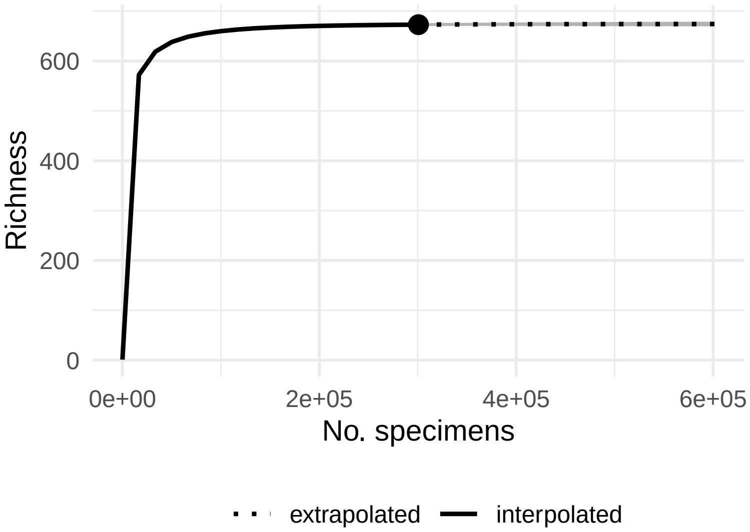


**Fig. S1**. Species collection curve for the ferns of Japan fit with the 'iNEXT' v.2.0.20 package (Hsieh et al., 2016). Grey shading indicates 95% confidence interval (not visible where it does not extend beyond the fitted line). Point indicates observed taxonomic richness (673 taxa, 95% confidence interval 670 to 676 taxa). Taxa include species, subspecies, and varieties, excluding hybrids. Sampling completeness is estimated to be 100% (95% confidence interval 100% to 100%).

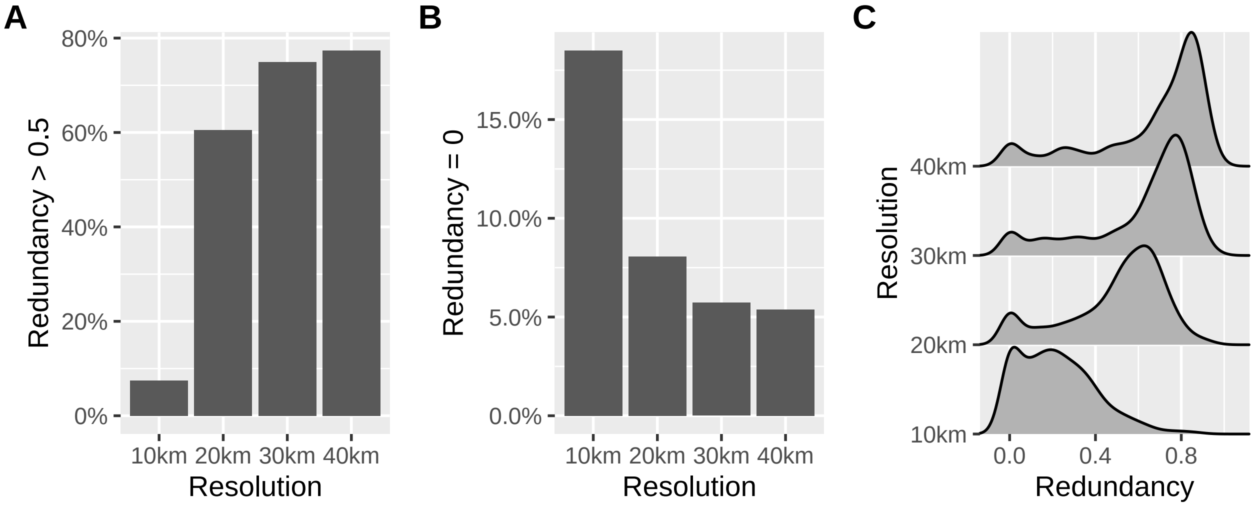


**Fig. S2**. Effect of grain size on sampling redundancy. (A) Percent of grid cells with redundancy > 0.5 by grain size (side length of square grid cells set to 10 km, 20 km, 30 km, or 40 km). (B) Percent of grid cells with redundancy = 0 by grain size. (C) Distribution of redundancy values by grain size.

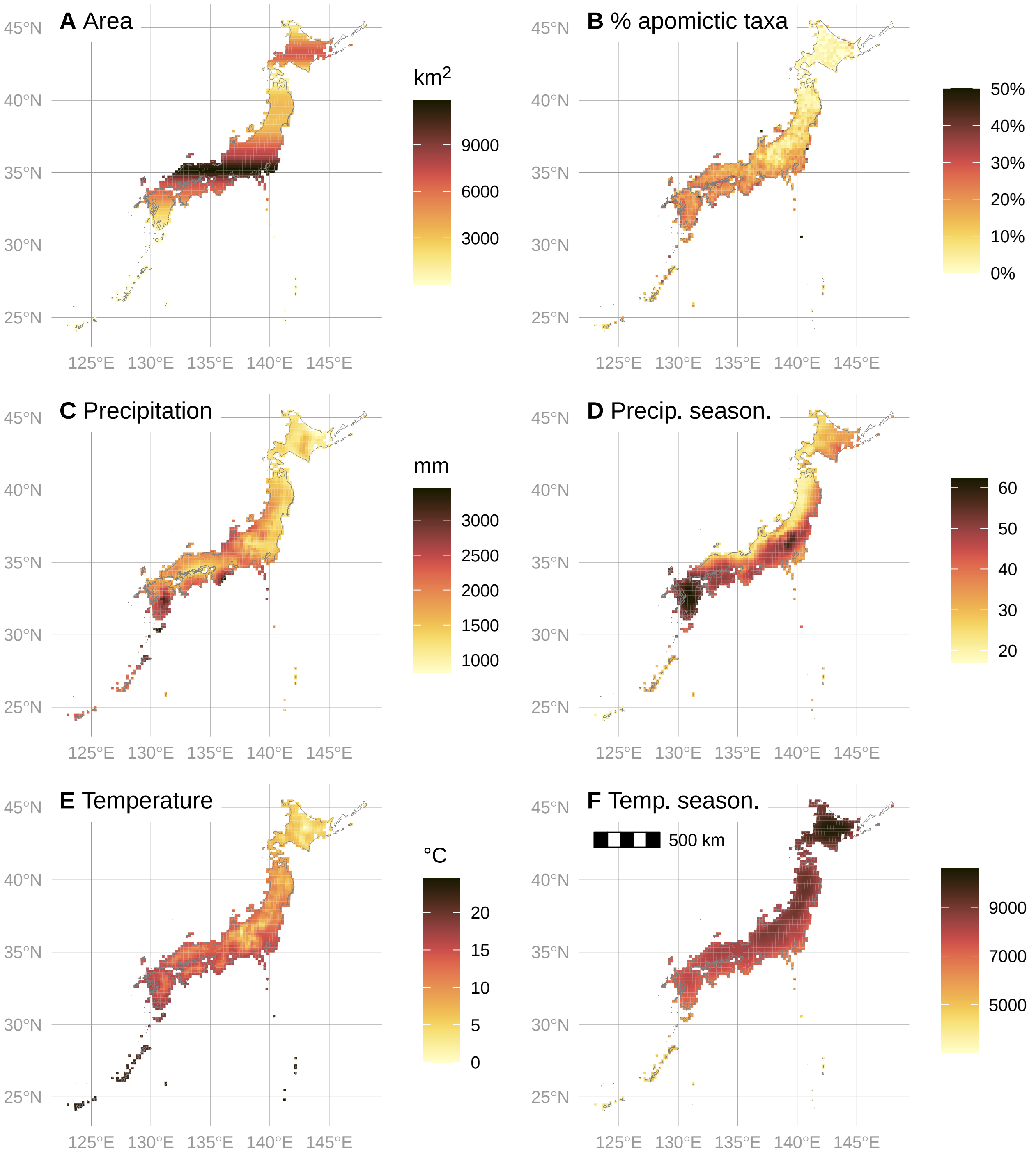


**Fig. S3**. Reproductive mode and environmental data in 20 km × 20 km grid cells across Japan. (A) rolling mean area (km$^2$) in 20 km latitudinal bands. (B) % apomictic taxa. (C) annual precipitation. (D) percipitation seasonality (coefficient of variation). (E) mean annual temperature. (F) temperature seasonality (standard deviation × 100). Data excluded from models not shown in this and subsequent plots unless otherwise mentioned.

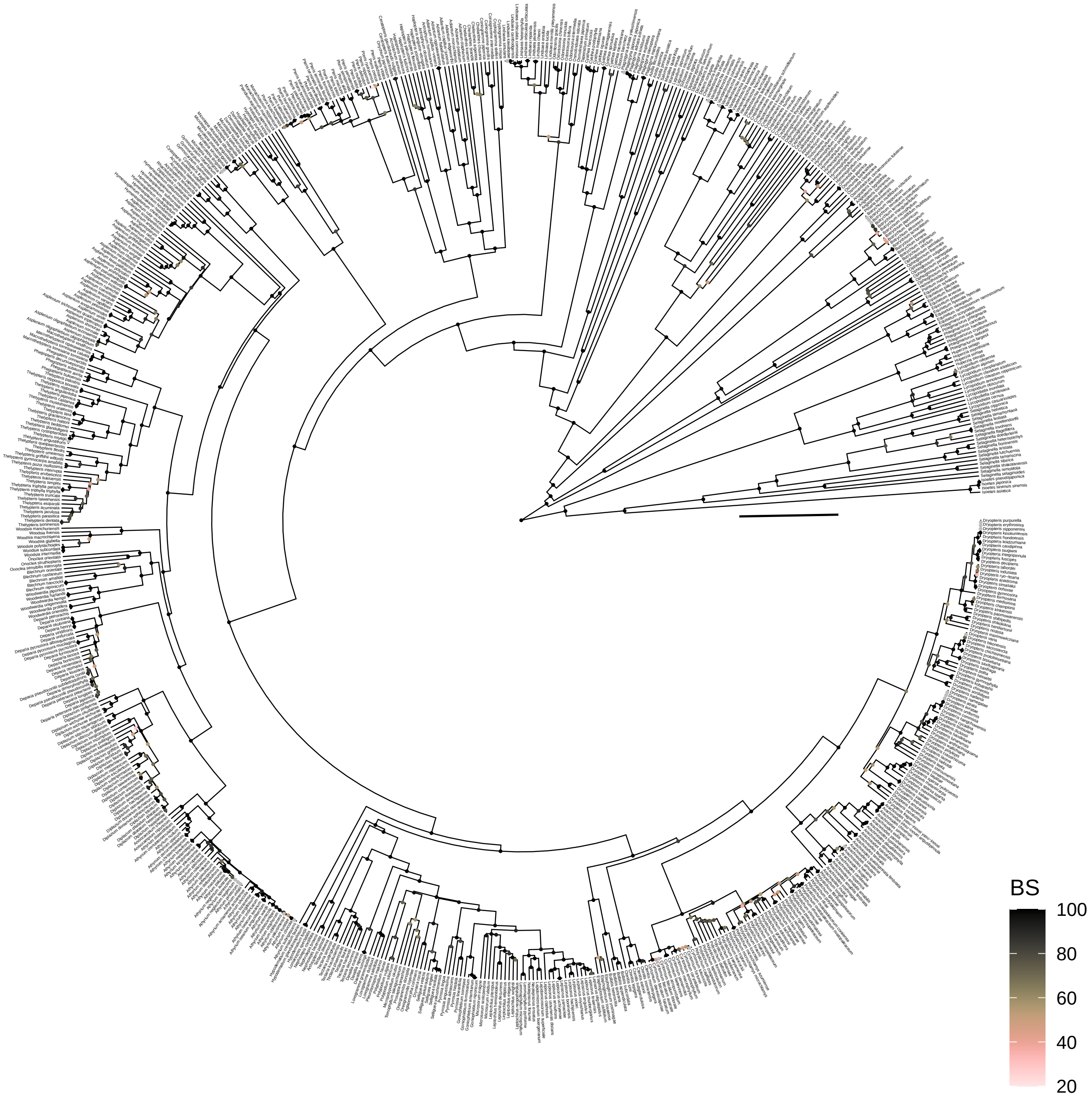


**Fig. S4**. Time tree of native Japanese ferns and lycophytes, excluding hybrids. Points at nodes indicate bootstrap support. Scalebar 100 my. Tree visualized with the 'ggtree' v.3.2.1 R package (Yu et al., 2017).

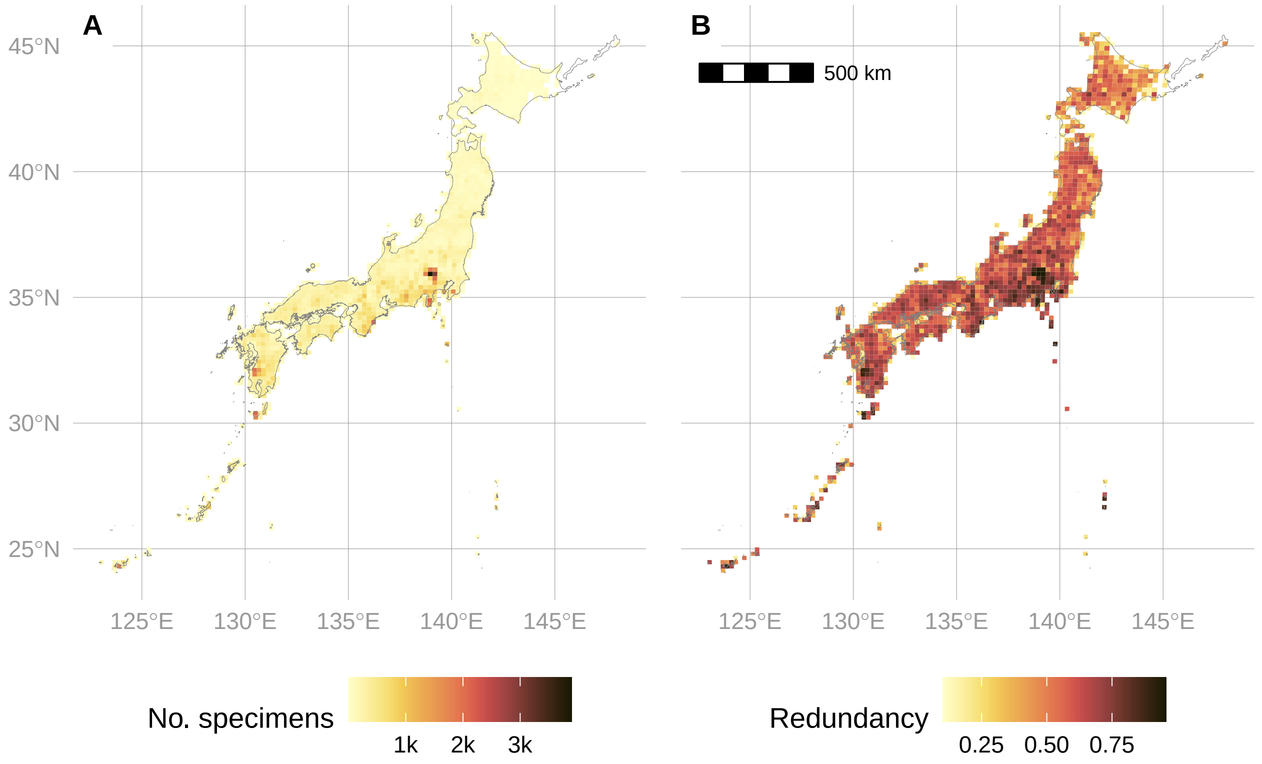


**Fig. S5**. Observed number of specimens (A) and sampling redundancy (B) per 20 km × 20 km grid cell in the ferns of Japan.

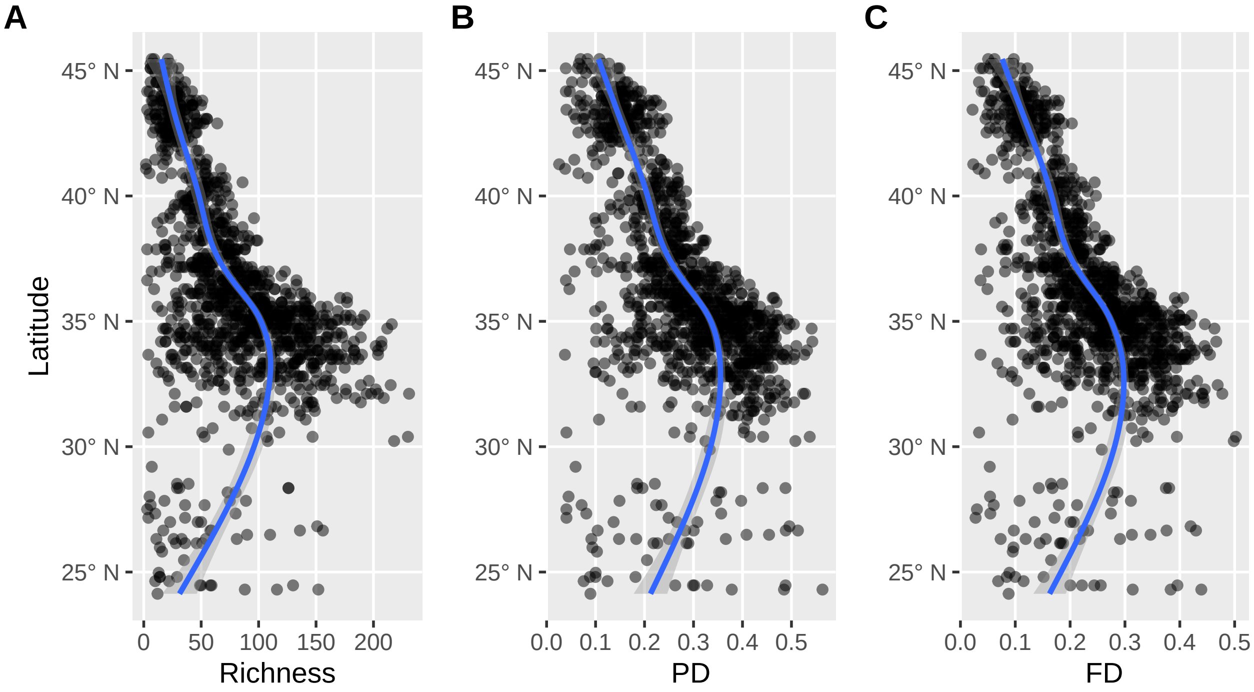


**Fig. S6**. (A) Raw taxonomic diversity, (B) phylogenetic diversity, and (C) functional diversity of the ferns of Japan plotted by latitude. For (B) and (C), raw branchlengths were transformed to relative values by dividing each branchlength by the sum of all branchlengths. Trendlines fit with general additive model using 'geom_smooth' function in the 'ggplot2' v.3.3.5.9000 R package (Wickham, 2016).

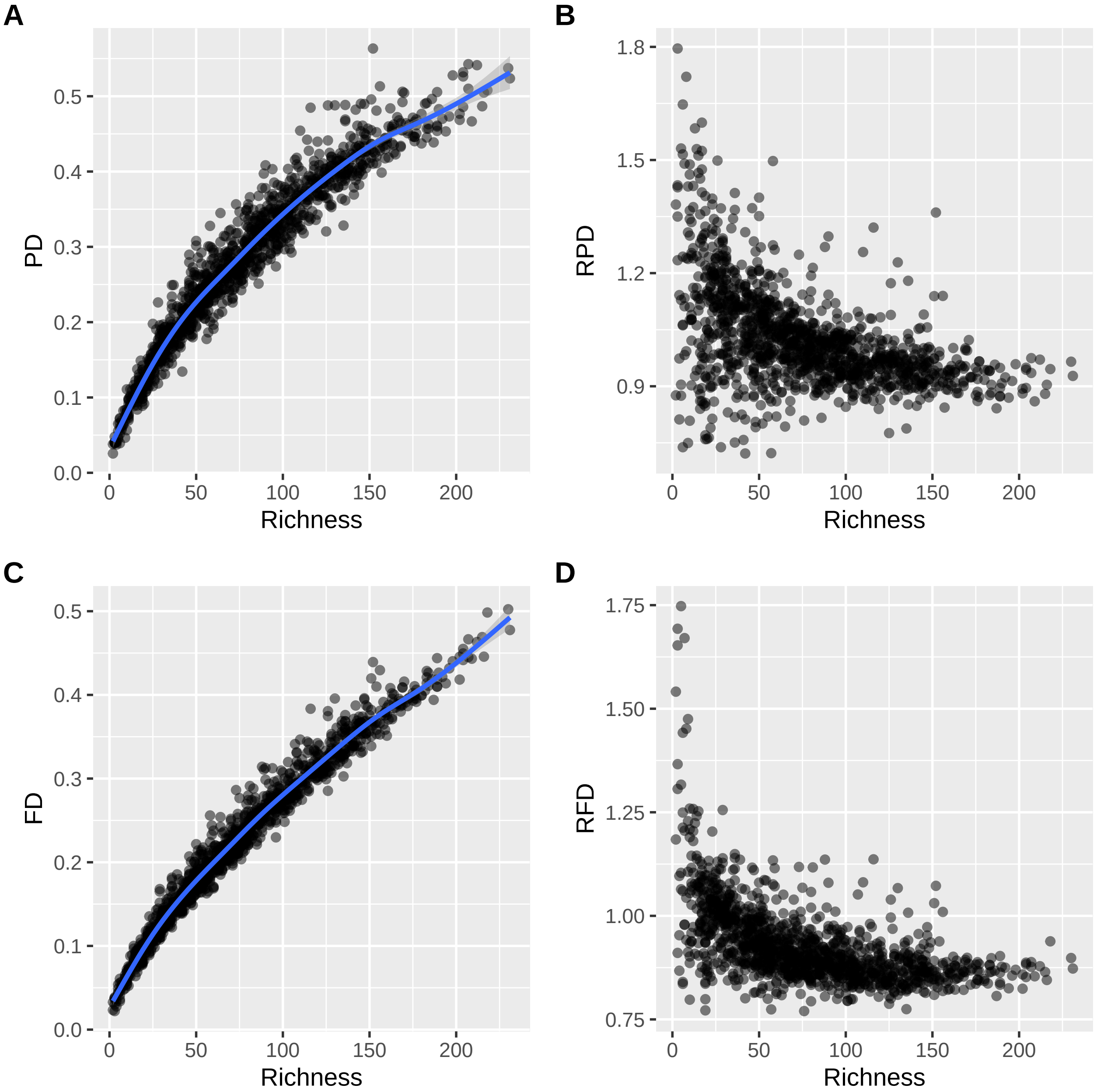


**Fig. S7**. Relationships between observed phylogenetic and functional diversity and taxonomic richness in the ferns of Japan. (A) Phylogenetic diversity (PD). (B) Relative phylogenetic diversity (RPD). (C) Functional diversity (FD). (D) Relative functional diversity (RFD). Trendlines in (A, C) fit with general additive model using 'geom_smooth' function in the 'ggplot2' v.3.3.5.9000 R package (Wickham, 2016).

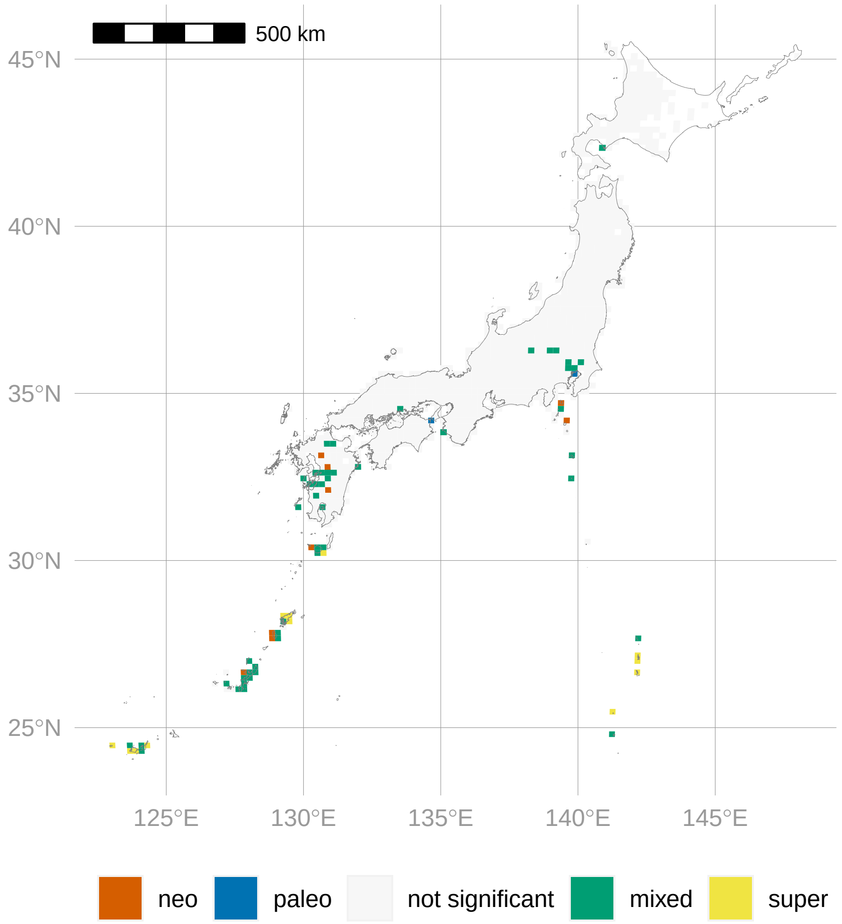


**Fig. S8**. Phylogenetic endemism of the ferns of Japan measured using CANAPE (categorical analysis of neo- and paleo-endemism), restricted dataset including only taxa endemic to Japan. ‘neo’ indicates neodendemic areas (those with an overabundance of short branches); ‘paleo’ indicates paleoendemic areas (those with an overabundance of long branches); ‘mixed’ indicates significantly endemic sites that have neither an overabundance of short nor long branches; ‘super’ indicates highly significantly endemic sites that have neither an overabundance of short nor long branches. For details, see Materials and Methods.

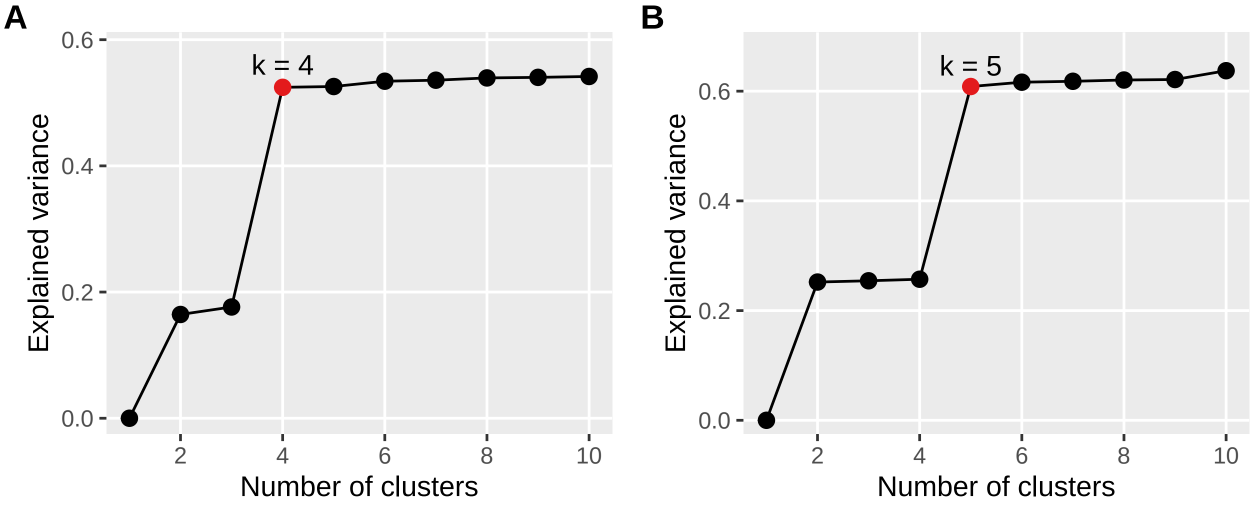


**Fig. S9**. Selection of *k* for bioregions. (A) taxonomic bioregions. (B) phylogenetic bioregions. Color indicates optimal *k* selected selected by the 'optimal_phyloregion' function in the R package 'phyloregion' v.1.0.6 (black, non-optimal; red, optimal).

**A** Taxonomic

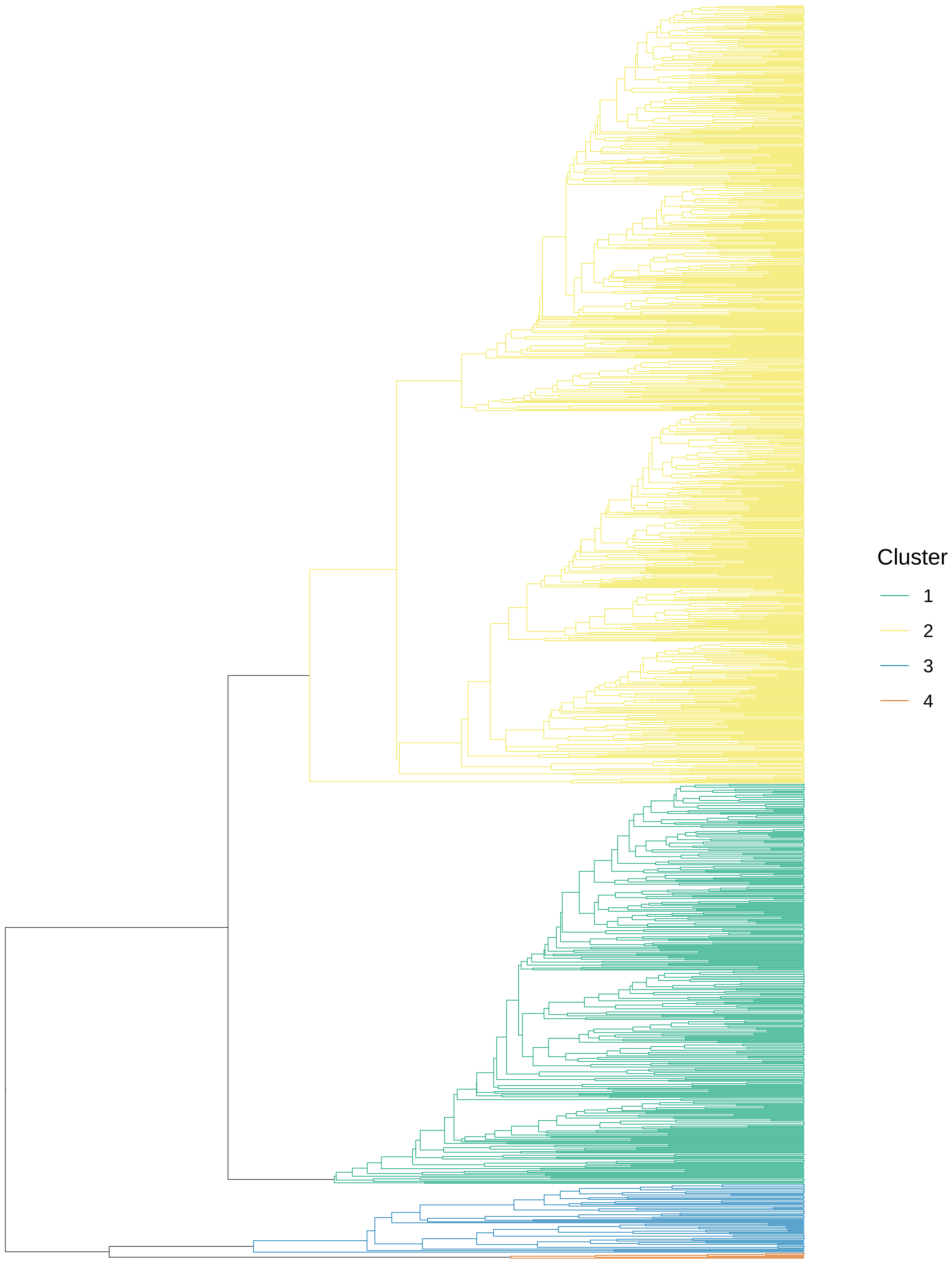


**Fig. S10**. Bioregion dendrograms. Tips are sites (names not shown). Dendrograms generated by clustering taxonomic (Sørensen) or phylogenetic (PhyloSor) distances between sites (see

Materials and Methods). Color shows bioregion; bioregions not consisting of more than two grid-cells each are lumped into the “Other” category. (A) taxonomic bioregions. (B) phylogenetic bioregions. Dendrograms visualized with the ‘ggtree’ v.3.2.1 R package (Yu et al., 2017).

**B** Phylogenetic

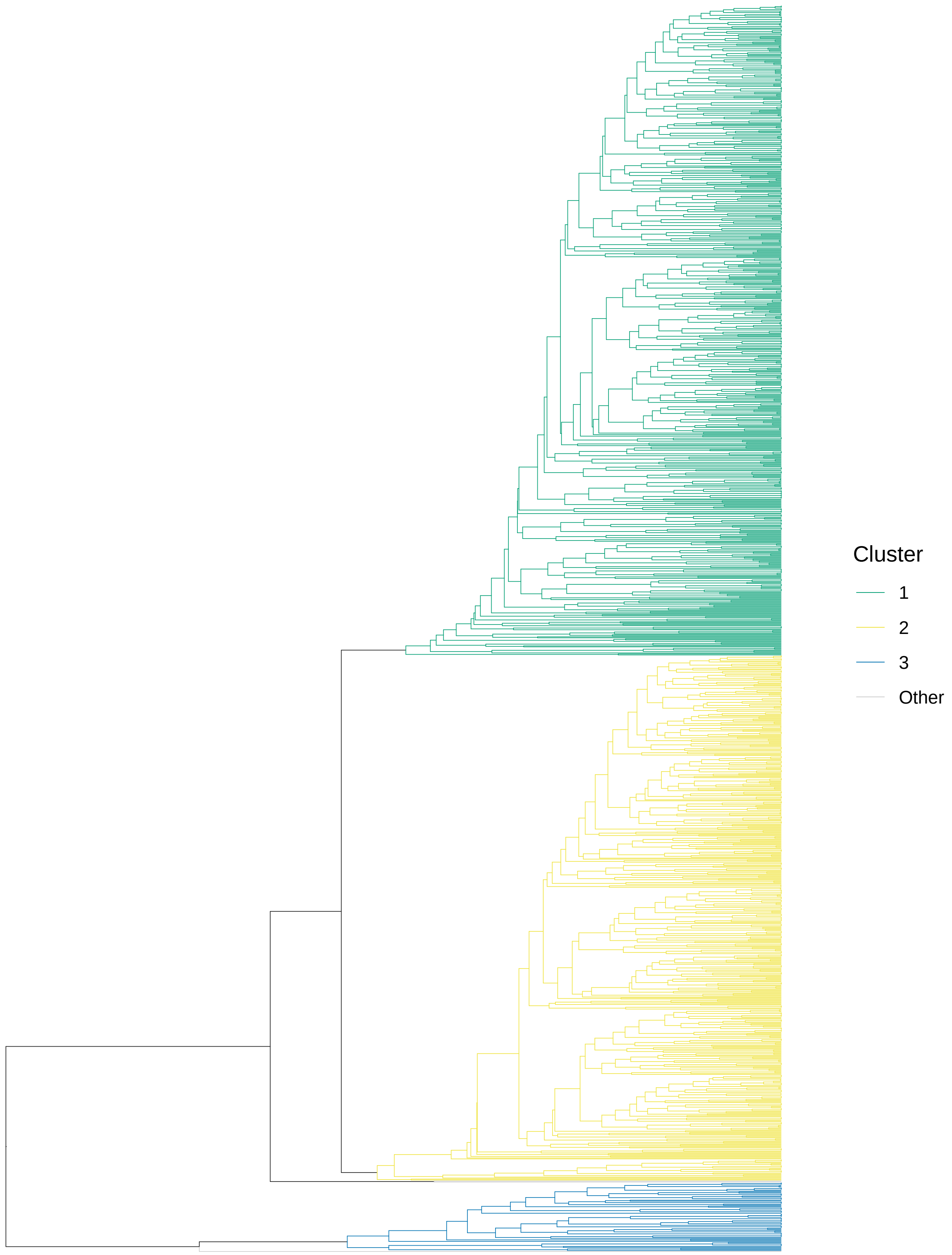


**Fig. S10**, continued.

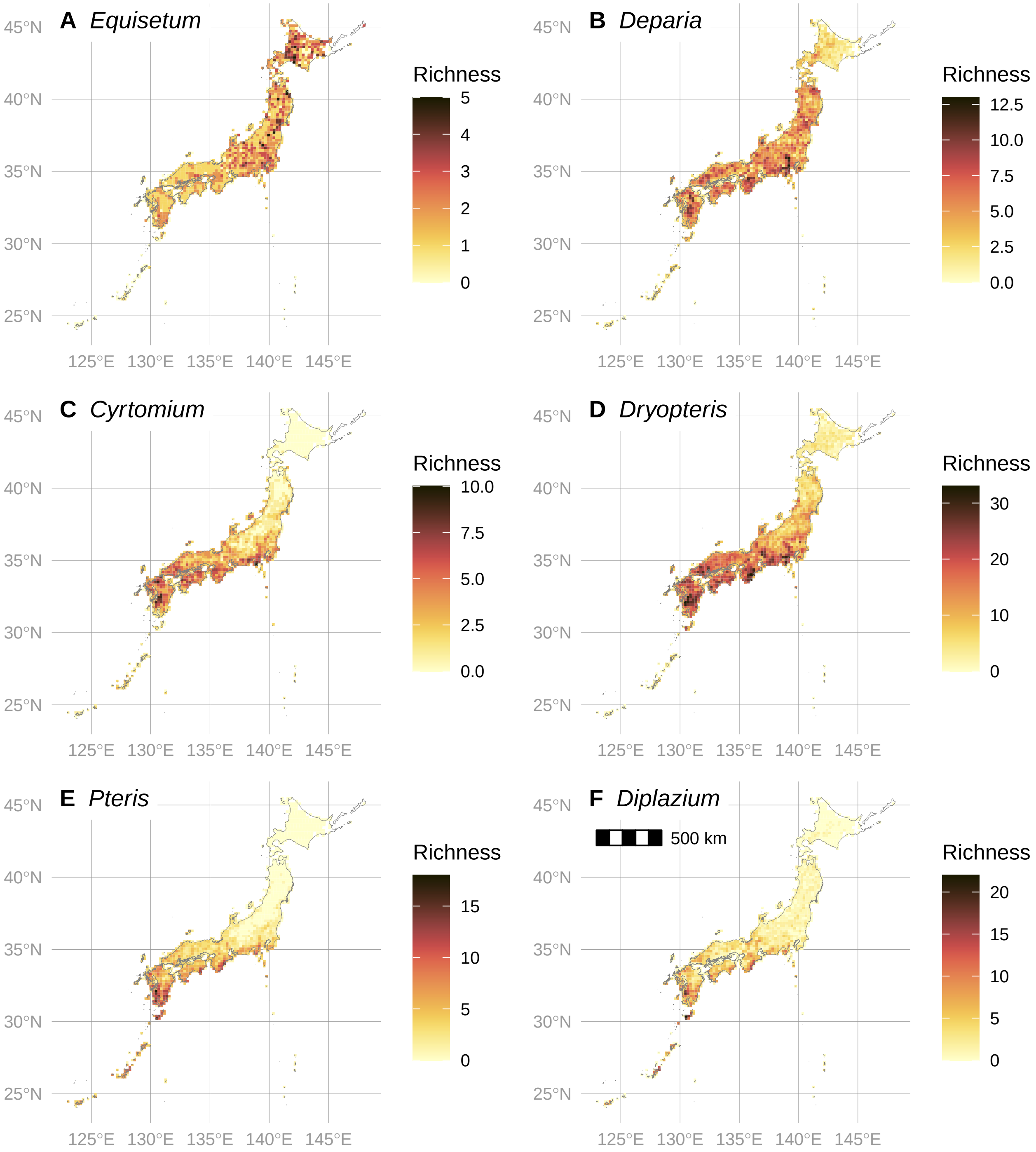


**Fig. S11**. Map of species richness for selected genera of ferns of Japan. (A) *Equisetum* is an example of a possible refugial lineage with greater species richness in northern Japan. (B) *Deparia* is an example of a genus with possible recent radiation in southern Honshu. (C)–(F) show genera with high rates of apomictic taxa (Fig. S12), which tend to be greatest in southern Honshu.

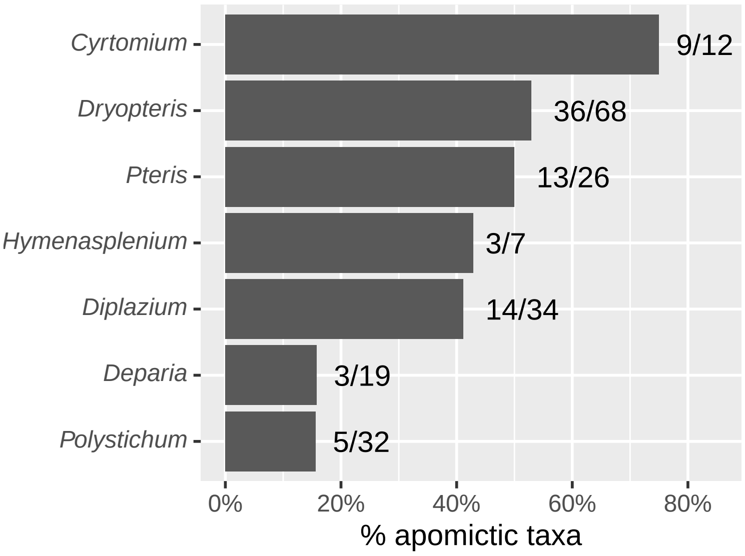


**Fig. S12**. Rate of apomixis in genera of native, non-hybrid Japanese ferns with > 1 apomictic taxon. Numbers next to each bar show number of apomictic taxa out of total richness per genus.

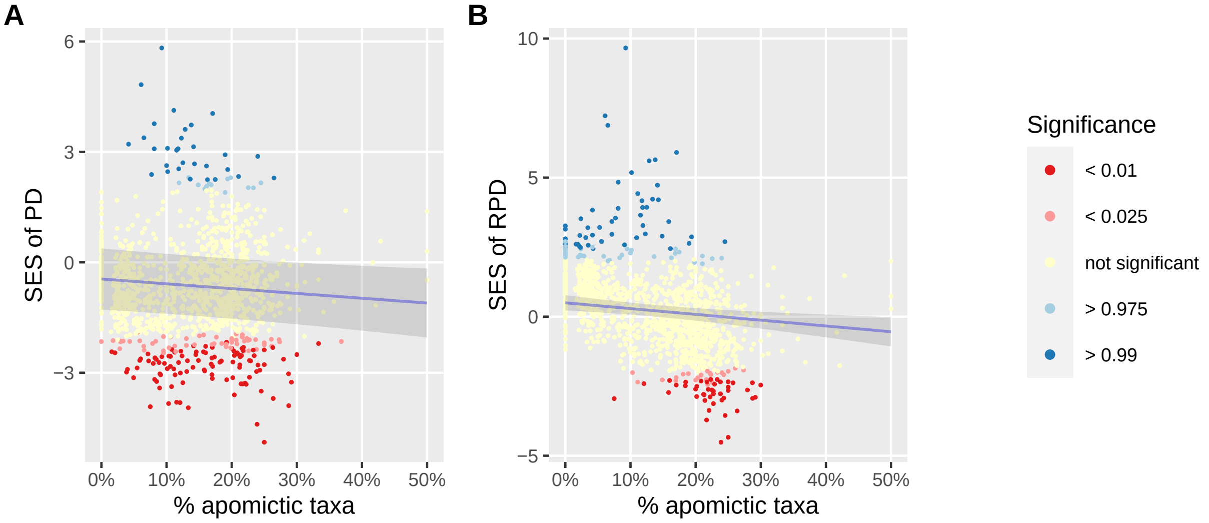


**Fig. S13**. Relationship between (A) phylogenetic diversity (PD) and (B) relative PD and % apomictic taxa, models inferred using phylogeny with branchlengths in units of expected genetic change (untransformed ML tree). Ribbon shows 95% confidence interval of model. Line fit with the focal predictor variable while averaging over other predictors. For RPD, the standard effect size (SES) was calculated by comparing raw values to a null distribution of 999 random values, and significance (*P*-value) calculated using a two-tailed test (see Materials and Methods).

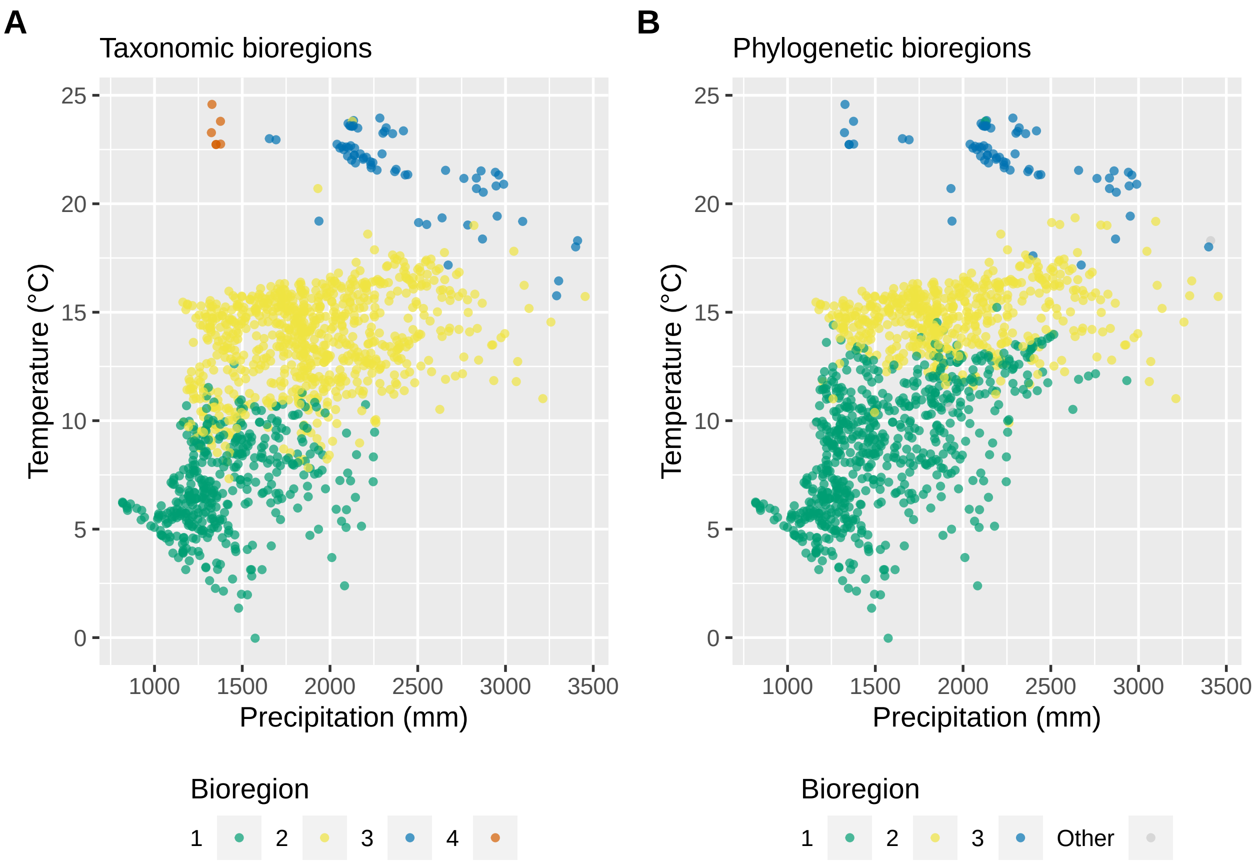


**Fig. S14**. Scatterplots of grid cell membership in (A) taxonomic or (B) phylogenetic bioregions arranged by mean annual temperature and annual precipitation. Points shown with partial transparency; darker areas indicate overlapping points. Bioregions not consisting of more than two grid-cells each are lumped into the "Other" category.

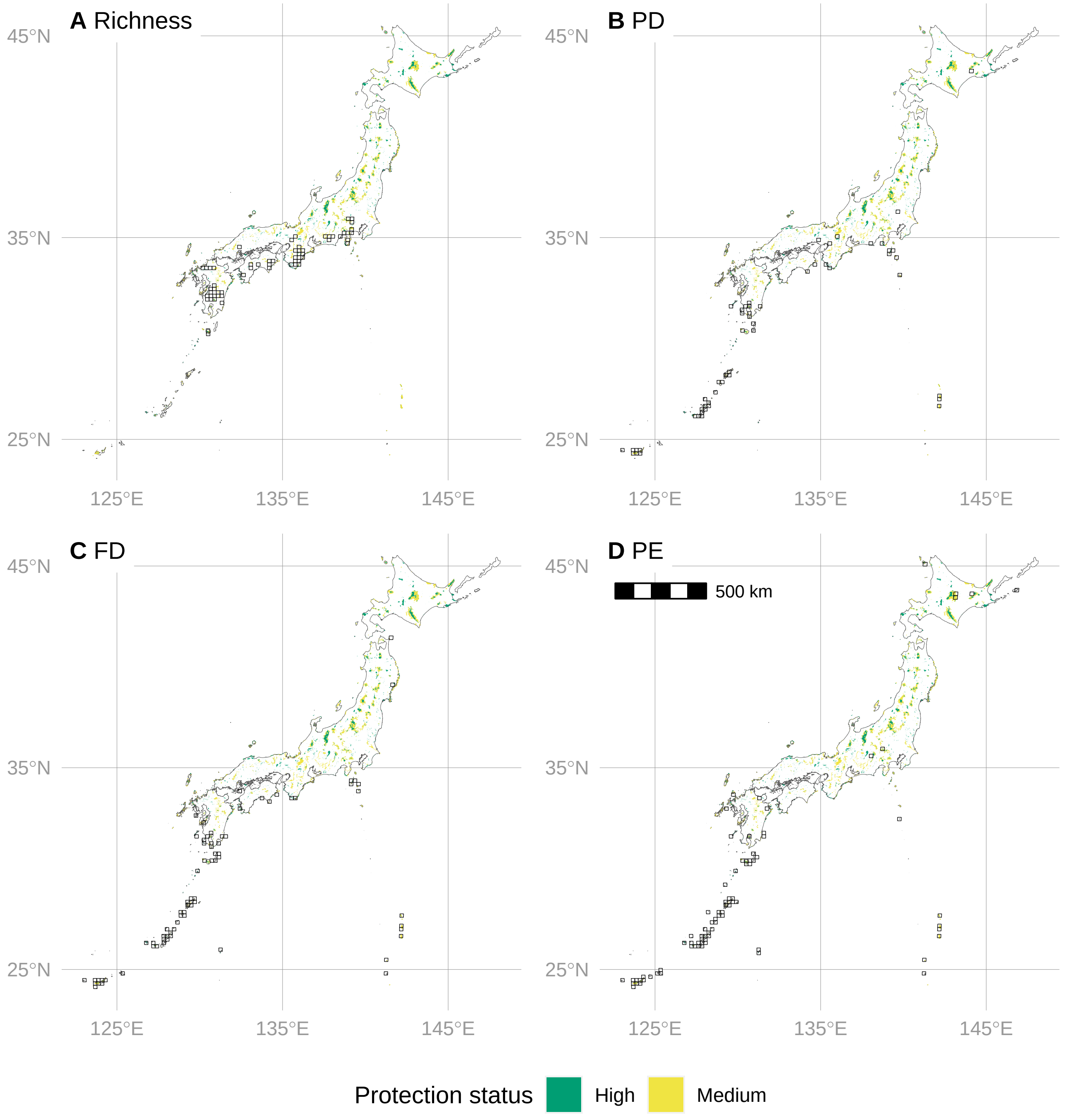


**Fig. S15**. Maps showing overlap of grid-cells with significantly high biodiversity for the ferns of Japan and protected areas. Grid-cells with black outlines indicate significantly high biodiversity. Biodiversity metrics include (A) taxon richness, (B) phylogenetic diversity (PD), (C) functional diversity (FD), and (D) phylogenetic endemism (PE). Significance of PD, FD, and PE assessed by a one-tailed test comparing observed values to a null distribution of 999 random values; for richness, grid-cells in the top 5% considered highly diverse (see Materials and Methods).

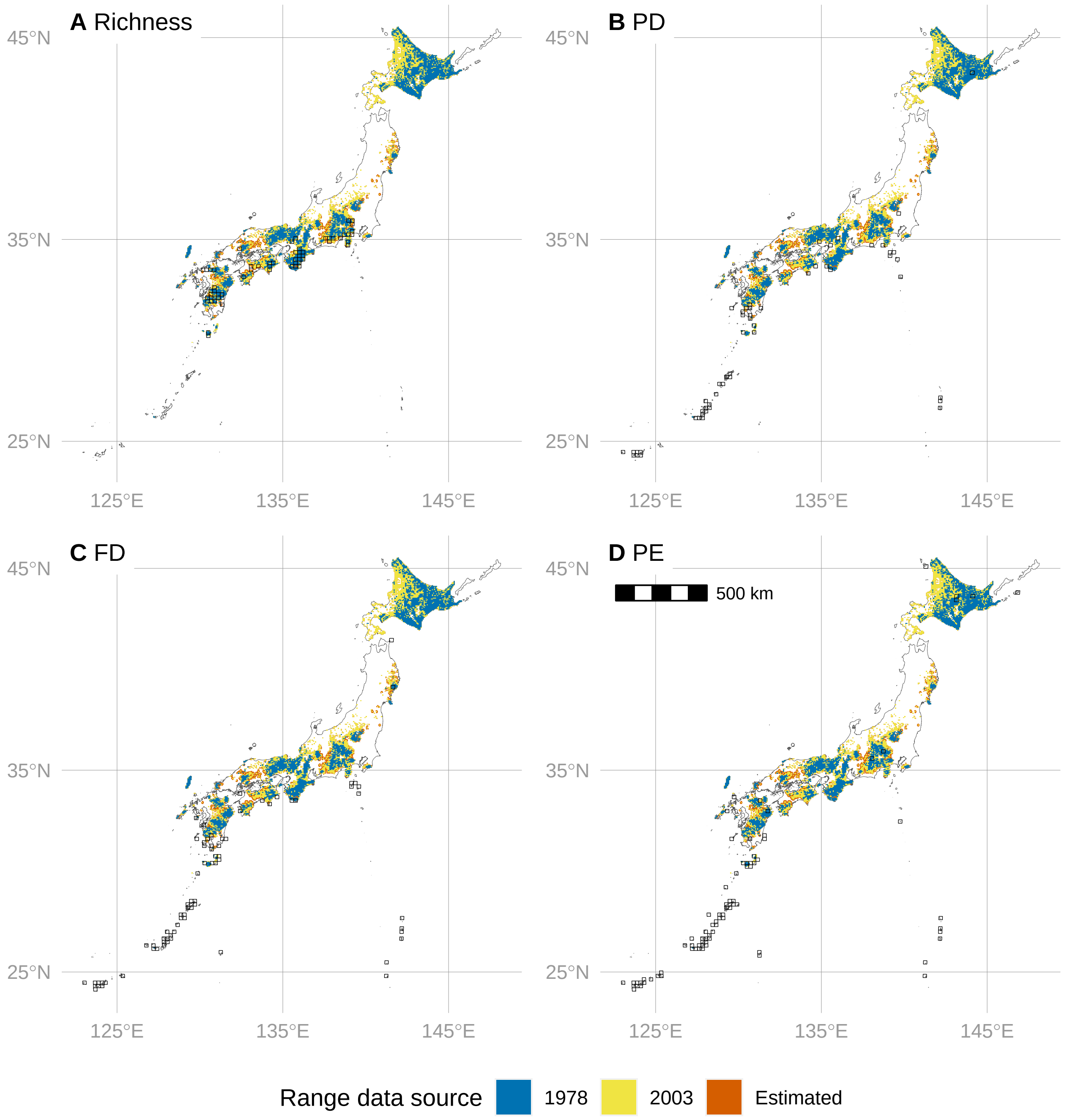


**Fig. S16**. Maps showing overlap of grid-cells with significantly high biodiversity for the ferns of Japan and distribution of Japanese deer (*Cervus nippon*). Grid-cells with black outlines indicate significantly high biodiversity. Biodiversity metrics include (A) taxon richness, (B) phylogenetic diversity (PD), (C) functional diversity (FD), and (D) phylogenetic endemism (PE). Significance of PD, FD, and PE assessed by a one-tailed test comparing observed values to a null distribution of 999 random values; for richness, grid-cells in the top 5% considered highly diverse (see Materials and Methods). Deer range data sources include 1978 survey, 2003 survey, and range estimated from model based on 2003 survey data. Deer ranges plotted in layers in ascending order (estimated, 2003, 1978) such that upper layers may obscure lower layers; however, deer ranges generally expanded from 1978 to estimated range, so very few sites are obscured.
