## Appendix S2 for "Spatial phylogenetics of Japanese ferns: Patterns, processes, and implications for conservation"

### Appendix S2: Data Exploration

Here we perform initial data exploration to determine the appropriate models to fit to the data.

All code is in R, and workflow managed with the `targets` package (https://github.com/ropensci/targets).

The code here is included in the repo at https://github.com/joelnitta/japan_ferns_spatial_phy.

#### Setup

Let's get started by loading packages and tidying our namespace:

```
# Load packages
library(tidyverse)
library(targets)
library(outliers)
library(ggforce)
library(assertr)
library(conflicted)

# Resolve possible namespace conflicts
conflict_prefer("map", "purrr")
conflict_prefer("select", "dplyr")
conflict_prefer("filter", "dplyr")
conflict_prefer("gather", "tidyr")
conflict_prefer("extract", "magrittr")
```

Also define a custom function we'll use later:

```
#' Run Grubb's test for outliers
#'
#' Similar to `outliers::grubbs.test()`, but returns the results in a tidy
#' dataframe
#'
#' @param x a numeric vector for data values.
#' @param type Integer value indicating test variant.
#' See `outliers::grubbs.test()`
#' @param ... Other arguments passed to `outliers::grubbs.test()`
#'
#' @return Tibble
grubbs_tidy <- function(x, type = 10, ...) {
  outliers::grubbs.test(x = x, type = type, ...) %>%
    broom::tidy() %>%
    dplyr::mutate(stat_name = c("g", "u")) %>%
```

```
    tidyr::pivot_wider(
      values_from = "statistic",
      names_from = "stat_name") %>%
    janitor::clean_names() %>%
    dplyr::select(g, u, p_value, method, alternative)
}
```

##### Load data

The dataset includes biodiversity metrics and environmental data measured on native, non-hybrid ferns of Japan.

All biodiversity metrics were calculated during the `targets` workflow (`_targets.R`), and are contained in the `biodiv_ferns_cent_figshare` object in the `targets` cache.

The next step uses `tar_load()` to load the dataset from the targets cache. Alternatively, this can be loaded from the data file as shown in the commented-out code below.

```
# Load the data from the targets workflow
tar_load(biodiv_ferns_cent_figshare)

# Alternatively, unzip `results.zip`
# (FIXME: add DOI when available)
# and read in the data from there
# biodiv_ferns_cent_figshare <- read_csv(here::here("japan_ferns_biodiv.csv"))
```

This dataframe actually has more variables than are needed for modeling, so we will subset it to only the relevant variables.

```
biodiv_ferns_cent_raw <-
  biodiv_ferns_cent_figshare %>%
  select(
    grids, lat, long, # grid-cell ID, latitude, longitude
    richness, pd_obs_z, fd_obs_z, rpd_obs_z, rfd_obs_z, # response vars
    percent_apo, temp, precip, precip_season, lat_area  # independent vars
  )
```

```
biodiv_ferns_cent_raw
```

```
## # A tibble: 1,239 x 13
##    grids   lat  long richness pd_obs_z fd_obs_z rpd_obs_z rfd_obs_z percent_apo
##    <chr> <dbl> <dbl>    <int>    <dbl>    <dbl>     <dbl>     <dbl>       <dbl>
##  1 4      24.1  124.       12   -0.269     1.67     0.645      1.96      0
##  2 114    24.3  124.      116    4.83      5.26     7.22       7.50      0.0610
##  3 115    24.3  124.      152    5.82      4.33     9.66       5.89      0.0926
##  4 116    24.3  124.       88    2.46      4.09     5.18       6.37      0.102
##  5 221    24.5  123.       59    1.69      3.87     3.52       4.71      0.0238
##  6 224    24.5  124.       58    3.38      5.08     6.88       5.02      0.0652
##  7 225    24.5  124.       50    3.08      3.71     4.84       3.33      0.0811
##  8 226    24.5  124.      130    3.73      4.15     5.64       5.50      0.138
```

```
##  9 227    24.5  124.        49    1.27     2.15      2.70       3.04      0.0556
## 10 337    24.6  124.        22   -0.879    0.826    -0.318      1.37      0
## # ... with 1,229 more rows, and 4 more variables: temp <dbl>, precip <dbl>,
## #   precip_season <dbl>, lat_area <dbl>
```

The columns are as follows:

- `grids`: ID code for each grid cell
- `lat`: latitude of grid cell centroid
- `long`: longitude of grid cell centroid
- `richness`: number of taxa
- `pd_obs_z`: standard effect size (SES) of phylogenetic diversity (pd)
- `fd_obs_z`: SES of functional diversity (fd)
- `rpd_obs_z`: SES of relative pd
- `rfd_obs_z`: SES of relative fd
- `percent_apo`: percent of apogamous taxa in that grid cell
- `temp`: mean annual temperature (10 x °C)
- `precip`: total annual precipitation (mm)
- `precip_season`: seasonality in precipitation
- `lat_area`: rolling mean of area in 0.2 degree latitudinal bands

##### Check for missing data

Check for missing data. If any rows are missing data, exclude them.

The next code chunk runs a logical check to see if any rows have missing data. It returns `TRUE` if there is no missing data, `FALSE` if there are any missing data:

```
biodiv_ferns_cent_raw %>%
  assert(
    not_na, everything(),
    error_fun = error_logical,
    success_fun = success_logical)
```

```
## [1] FALSE
```

At least one row has missing data. Let's see what variables are missing.

```
# Get the row numbers of rows with missing data
error_rows <-
  biodiv_ferns_cent_raw %>%
  assert(not_na, everything(), error_fun = error_df_return) %>%
  pull(index) %>%
  unique()

# Inspect the rows with missing data
biodiv_ferns_cent_raw %>%
  slice(error_rows) %>%
  select(grids, richness, percent_apo, temp, precip, precip_season)
```

```
## # A tibble: 3 x 6
```

```
##   grids richness percent_apo  temp precip precip_season
##   <chr>     <int>       <dbl> <dbl>  <dbl>         <dbl>
## 1 4438          1      NA       192   1937            50
## 2 994           4       1       NaN    NaN           NaN
## 3 1103         48       0.0882  NaN    NaN           NaN
```

The missing data are due to two grid-cells that lack environmental data from the WorldClim dataset and one grid-cell that only had a single species with no breeding-system data. Let's remove these, then continue.

```
# Exclude rows with missing data
biodiv_ferns_cent_no_missing <-
  biodiv_ferns_cent_raw %>%
  slice(-error_rows)

# Double check that the resulting dataframe has no missing data
assert(biodiv_ferns_cent_no_missing,
       not_na, everything(),
       success_fun = success_logical)
```

```
## [1] TRUE
```

##### Check for outliers

We will first check for outliers in the data by visualizing box plots. Dots at the extremes may be outliers.

```
biodiv_ferns_cent_no_missing %>%
  select(-lat, -long) %>%
  pivot_longer(-grids, names_to = "variable") %>%
  ggplot(aes(x = variable, y = value)) +
  geom_boxplot() +
  facet_wrap(vars(variable), scales = "free") +
  theme(
    axis.title.x = element_blank(),
    axis.text.x = element_blank()
  )
```

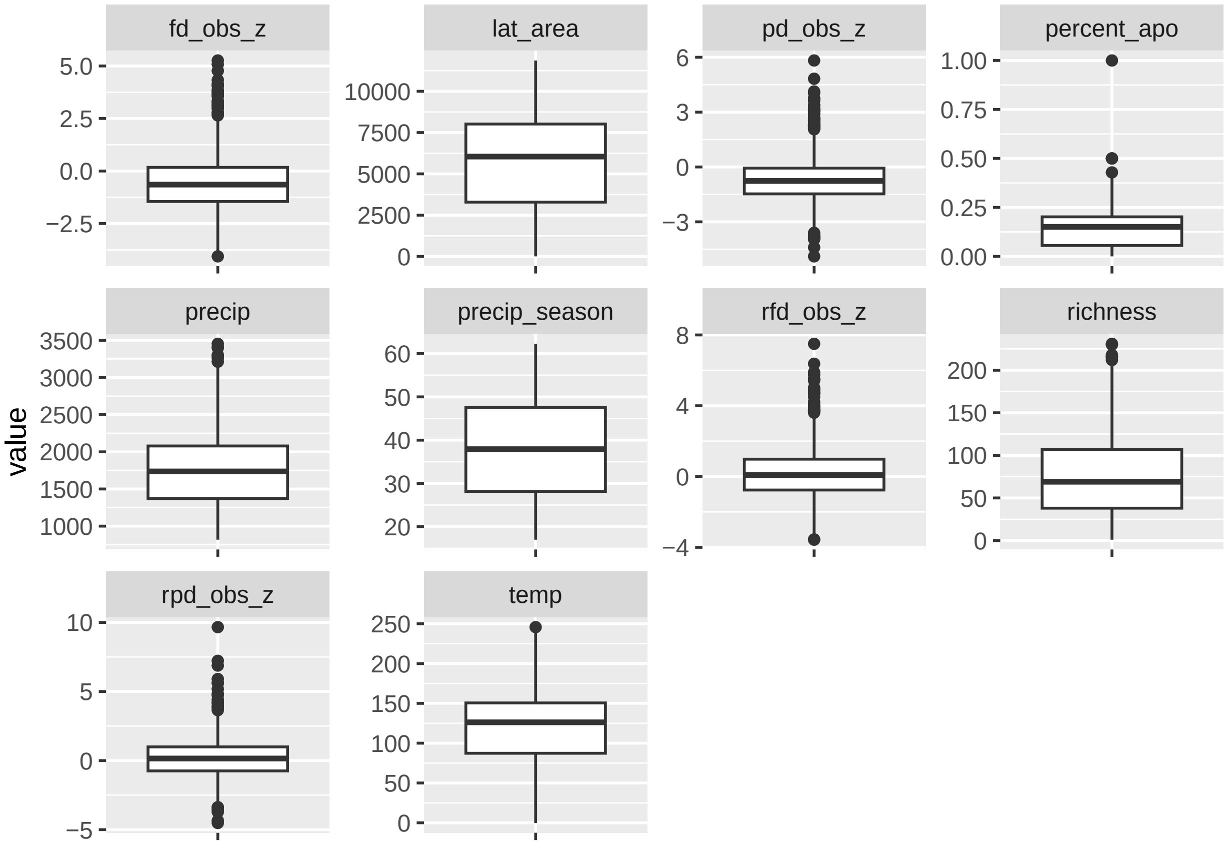


There are a few points beyond the whiskers that may be outliers. In particular, there is one value of `percent_apo` that seems quite outside the normal range.

Next, check for outliers using Grubb's test. Significant *p*-values indicate that the most extreme value is an outlier.

```
biodiv_ferns_cent_no_missing %>%
  select(-lat, -long) %>%
  pivot_longer(-grids, names_to = "variable") %>%
  group_by(variable) %>%
  nest() %>%
  mutate(map_df(data, ~grubbs_tidy(.$value))) %>%
  select(-data) %>%
  ungroup() %>%
  select(variable:p_value) %>%
  mutate(signif = p_value < 0.05)
```

```
## # A tibble: 10 x 5
##    variable          g     u        p_value signif
##    <chr>         <dbl> <dbl>          <dbl> <lgl>
##  1 richness       3.28 0.991 0.619          FALSE
##  2 pd_obs_z       5.14 0.979 0.000148       TRUE
##  3 fd_obs_z       4.38 0.984 0.00677        TRUE
##  4 rpd_obs_z      6.78 0.963 0.00000000494 TRUE
##  5 rfd_obs_z      5.36 0.977 0.0000430      TRUE
```

```
##  6 percent_apo    9.67 0.924 0            TRUE
##  7 temp           2.82 0.994 1            FALSE
##  8 precip         3.60 0.989 0.189        FALSE
##  9 precip_season  2.17 0.996 1            FALSE
## 10 lat_area       1.85 0.997 1            FALSE
```

Grubb's test confirms that the percent apomictic value is an outlier. It also indicates that the highest values of the variables `fd_obs_z`, `pd_obs_z`, `rfd_obs_z`, and `rpd_obs_z` may be outliers.

One reason for artificially extreme values could be undersampling of species. Let's check richness of these possible outlier grid-cells.

```
biodiv_ferns_cent_no_missing %>%
  slice_max(n = 1, order_by = percent_apo, with_ties = TRUE) %>%
  select(grids:long, percent_apo, richness)
```

```
## # A tibble: 1 x 5
##   grids   lat  long percent_apo richness
##   <chr> <dbl> <dbl>       <dbl>    <int>
## 1 343    24.6  126.           1        1
```

There is just one species in the grid-cell that obviously contributes to the 100% apomixis rate. This data point should be excluded.

What about the other possible outliers?

```
biodiv_ferns_cent_no_missing %>%
  slice_max(n = 2, order_by = fd_obs_z) %>%
  select(grids:long, fd_obs_z, richness)
```

```
## # A tibble: 2 x 5
##   grids   lat  long fd_obs_z richness
##   <chr> <dbl> <dbl>    <dbl>    <int>
## 1 114    24.3  124.     5.26      116
## 2 224    24.5  124.     5.08       58
```

```
biodiv_ferns_cent_no_missing %>%
  slice_max(n = 2, order_by = pd_obs_z) %>%
  select(grids:long, pd_obs_z, richness)
```

```
## # A tibble: 2 x 5
##   grids   lat  long pd_obs_z richness
##   <chr> <dbl> <dbl>    <dbl>    <int>
## 1 115    24.3  124.     5.82      152
## 2 114    24.3  124.     4.83      116
```

```
biodiv_ferns_cent_no_missing %>%
  slice_max(n = 2, order_by = rfd_obs_z) %>%
  select(grids:long, rfd_obs_z, richness)
```

```
## # A tibble: 2 x 5
##   grids   lat  long rfd_obs_z richness
##   <chr> <dbl> <dbl>     <dbl>    <int>
```

```
## 1 114    24.3  124.       7.50       116
## 2 116    24.3  124.       6.37        88
```

```
biodiv_ferns_cent_no_missing %>%
  slice_max(n = 2, order_by = rpd_obs_z) %>%
  select(grids:long, rpd_obs_z, richness)
```

```
## # A tibble: 2 x 5
##   grids   lat  long rpd_obs_z richness
##   <chr> <dbl> <dbl>     <dbl>    <int>
## 1 115    24.3  124.      9.66      152
## 2 114    24.3  124.      7.22      116
```

None of these have an extremely low number of species.

So we will conservatively only remove the grid-cell with unusually high % apomictic taxa, then run Grubb's test again.

```
biodiv_ferns_cent_no_outliers <-
  biodiv_ferns_cent_no_missing %>%
  filter(percent_apo != max(percent_apo))

biodiv_ferns_cent_no_outliers %>%
  pull(percent_apo) %>%
  grubbs_tidy()
```

```
## # A tibble: 1 x 5
##       g     u p_value method                      alternative
##   <dbl> <dbl>   <dbl> <chr>                       <chr>
## 1  4.23 0.985  0.0136 Grubbs test for one outlier highest value 0.5 is an outli~
```

OK, no more outliers in `percent_apo`.

##### Check for normality

Check for normality with histograms and QQ-plots after removing missing data and outliers.

First inspect histograms:

```
biodiv_ferns_cent_no_outliers %>%
  select(-lat, -long) %>%
  pivot_longer(-grids, names_to = "variable") %>%
  ggplot(aes(x = value)) +
  geom_histogram(bins = 30) +
  facet_wrap(vars(variable), scales = "free")
```

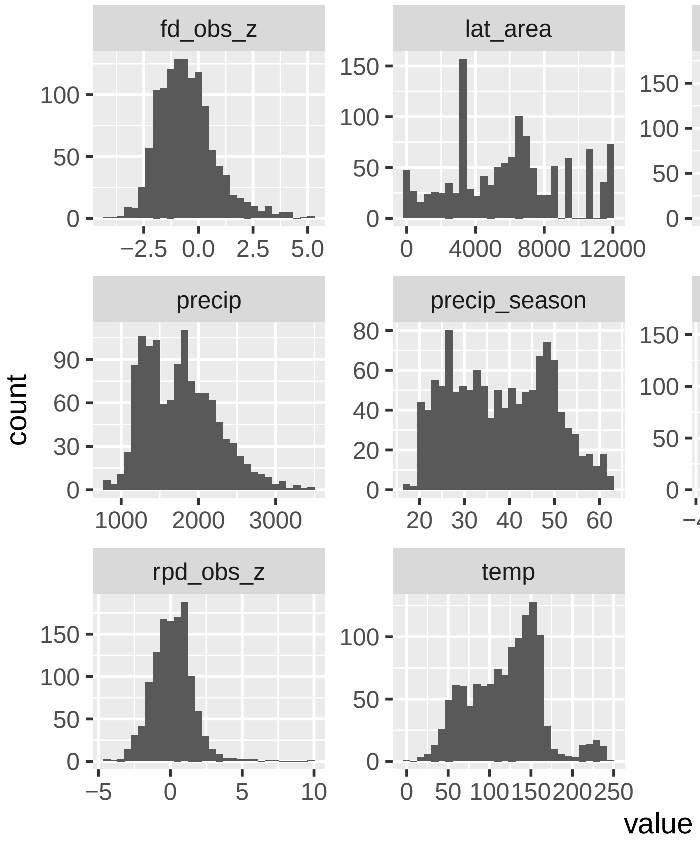


Next inspect qq-plots:

```
biodiv_ferns_cent_no_outliers %>%
  select(-lat, -long) %>%
  pivot_longer(-grids, names_to = "variable") %>%
  ggplot(aes(sample = value)) +
  stat_qq() +
  stat_qq_line() +
  facet_wrap(~variable, scales = "free")
```

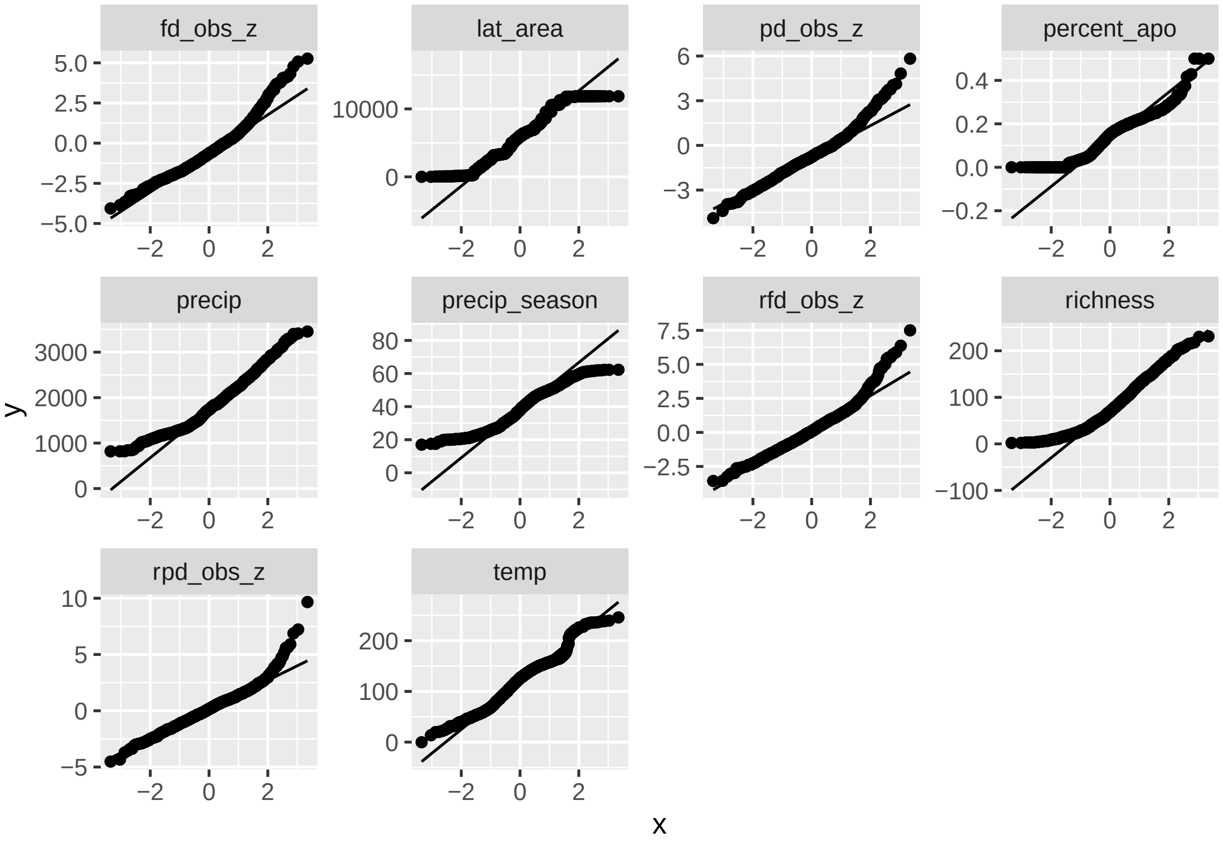


The response variables `fd_obs_z`, `pd_obs_z`, `rfd_obs_z`, `rpd_obs_z` all look decent. `richness` is skewed, but it is count data and we aren't using a Gaussian model for that anyways (poisson or negative binomial), so that's OK.

##### Check for zeros in count data

Zero-inflated data can be a problem for linear models.

```
biodiv_ferns_cent_no_outliers %>%
  mutate(rich_zero = richness == 0) %>%
  arrange(richness) %>%
  summarize(
    total_zero = sum(rich_zero),
    perc_zero = sum(rich_zero) / nrow(.))
```

```
## # A tibble: 1 x 2
##   total_zero perc_zero
##        <int>     <dbl>
## 1          0         0
```

No zeros in our count data (richness).

##### Plot raw data

Here we plot response variables (row labels) vs. indep variables (column labels) and add a trend line for each to get an idea of the possible model fits.

```
ggplot(biodiv_ferns_cent_no_outliers, aes(x = .panel_x, y = .panel_y)) +
  geom_point(alpha = 0.5, shape = 16, size = 0.5) +
  geom_smooth() +
  facet_matrix(
    cols = vars(temp, precip, precip_season, percent_apo),
    rows = vars(richness, pd_obs_z, fd_obs_z, rpd_obs_z, rfd_obs_z))
```

```
## Warning in rows == cols: longer object length is not a multiple of shorter
## object length

## `geom_smooth()` using method = 'gam' and formula 'y ~ s(x, bs = "cs")'
```

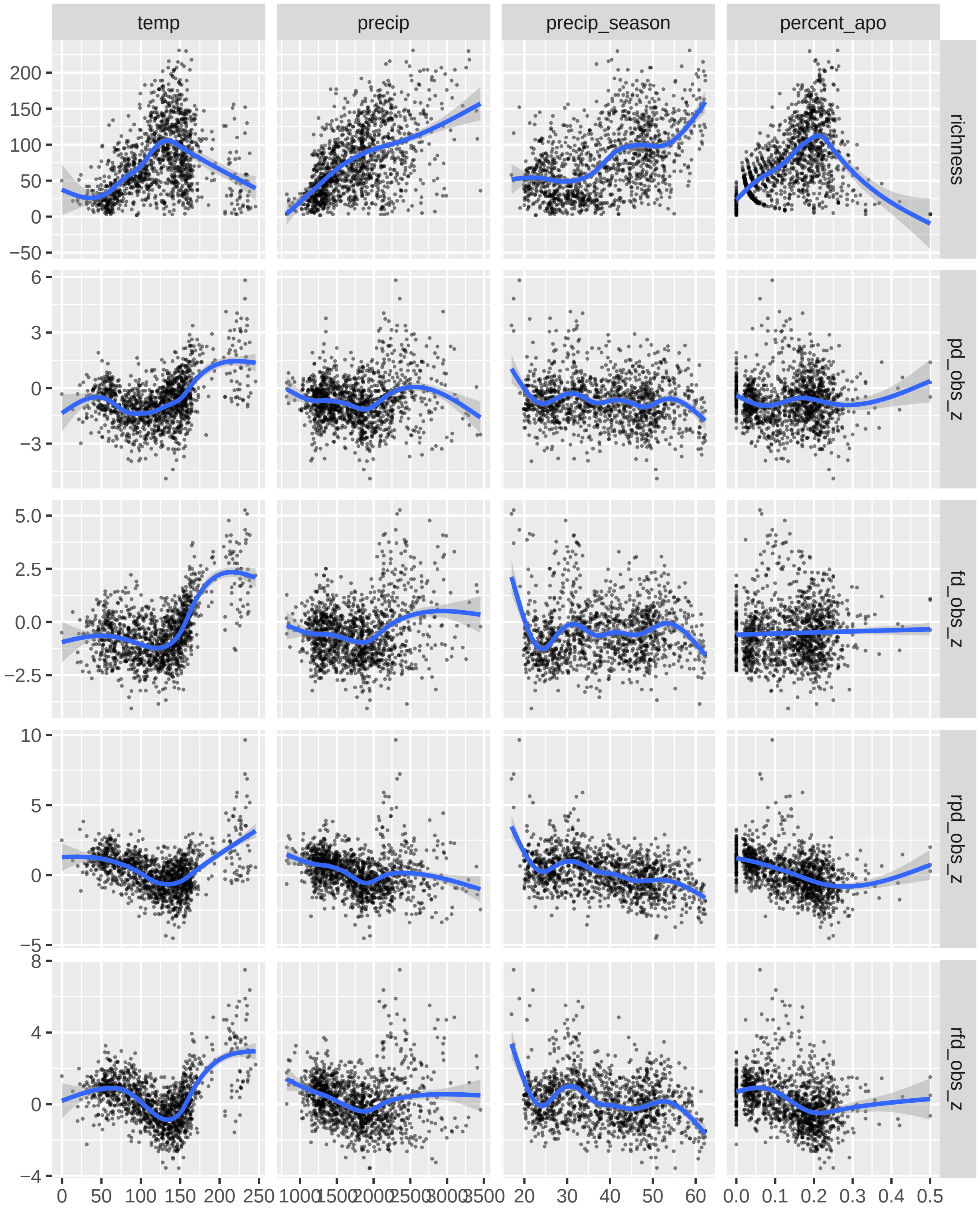


`richness` vs. `temp` definitely shows a hump-shaped trend line, and the other response variables vs. `temp` also show one if not quite as strongly. The other relationships appear to be linear or uncorrelated (the bend in the line for `percent_apo` vs. `richness` seems to be due to a relatively few number of points with > 30% apomictic taxa, but otherwise it looks linear).

##### Check for overdispersion in richness

Since richness is count data (only takes positive values), we will use either a poisson or negative binomial to model its distribution. The poisson distribution requires that the ratio of residual variance to degrees of freedom be approximately 1; significantly different ratios are "overdispersed" or "underdispersed". Check for this with a dispersion test.

```
# Create general linear model with poisson distribution
richness_poisson_mod <- glm(
  richness ~ temp + I(temp^2) + precip + precip_season,
  data = biodiv_ferns_cent_no_outliers, family = poisson)

# Check ratio of residual variance to df
# (should be near 1 to meet poisson assumptions)
richness_poisson_mod$deviance / richness_poisson_mod$df.residual
```

```
## [1] 14.77217
```

That's very overdispersed. So we won't use poisson for `richness`, but negative binomial instead.

##### Session information

```
sessionInfo()
```

```
## R version 4.1.2 (2021-11-01)
## Platform: x86_64-pc-linux-gnu (64-bit)
## Running under: Ubuntu 20.04.3 LTS
##
## Matrix products: default
## BLAS/LAPACK: /usr/lib/x86_64-linux-gnu/openblas-pthread/libopenblasp-r0.3.8.so
##
## locale:
##  [1] LC_CTYPE=en_US.UTF-8       LC_NUMERIC=C
##  [3] LC_TIME=en_US.UTF-8        LC_COLLATE=en_US.UTF-8
##  [5] LC_MONETARY=en_US.UTF-8    LC_MESSAGES=C
##  [7] LC_PAPER=en_US.UTF-8       LC_NAME=C
##  [9] LC_ADDRESS=C               LC_TELEPHONE=C
## [11] LC_MEASUREMENT=en_US.UTF-8 LC_IDENTIFICATION=C
##
## attached base packages:
## [1] stats     graphics  grDevices datasets  utils     methods   base
##
## other attached packages:
##  [1] ggforce_0.3.3      outliers_0.14      forcats_0.5.1      stringr_1.4.0
##  [5] dplyr_1.0.7        purrr_0.3.4        readr_2.1.1        tidyr_1.1.4
##  [9] tibble_3.1.6       ggplot2_3.3.5.9000 tidyverse_1.3.1    rgnparser_0.2.0
## [13] canaper_0.0.2      testthat_3.1.1     future.callr_0.7.0 future_1.23.0
## [17] spaMM_3.9.25       cachem_1.0.6       memoise_2.0.1      ggridges_0.5.3
## [21] ggtext_0.1.1       RColorBrewer_1.1-2 patchwork_1.1.1    glue_1.6.0
```

```
## [25] clustermq_0.8.95.2 scico_1.3.0        phyloregion_1.0.6  sf_1.0-5
## [29] magrittr_2.0.1     assertr_2.8        broom_0.7.10       jntools_0.1.0
## [33] picante_1.8.2      nlme_3.1-153       maps_3.4.0         janitor_2.1.0
## [37] readxl_1.3.1       assertthat_0.2.1   here_1.0.1         FD_1.0-12
## [41] vegan_2.5-7        lattice_0.20-45    permute_0.9-5      geometry_0.4.5
## [45] ape_5.6            ade4_1.7-18        checkr_0.5.0       conflicted_1.1.0
## [49] tarchetypes_0.4.0  targets_0.9.0
##
## loaded via a namespace (and not attached):
##   [1] snow_0.4-4          backports_1.4.1     fastmatch_1.1-3
##   [4] plyr_1.8.6          igraph_1.2.10       sp_1.4-6
##   [7] splines_4.1.2       listenv_0.8.0       digest_0.6.29
##  [10] htmltools_0.5.2     foreach_1.5.1       fansi_0.5.0
##  [13] cluster_2.1.2       tzdb_0.2.0          globals_0.14.0
##  [16] modelr_0.1.8        clustMixType_0.2-15 colorspace_2.0-2
##  [19] rvest_1.0.2         haven_2.4.3         xfun_0.29
##  [22] callr_3.7.0         crayon_1.4.2        jsonlite_1.7.2
##  [25] phangorn_2.8.1      iterators_1.0.13    polyclip_1.10-0
##  [28] registry_0.5-1      gtable_0.3.0        abind_1.4-5
##  [31] scales_1.1.1        DBI_1.1.2           Rcpp_1.0.7
##  [34] gridtext_0.1.4      magic_1.5-9         units_0.7-2
##  [37] proxy_0.4-26        httr_1.4.2          dismo_1.3-5
##  [40] betapart_1.5.4      ellipsis_0.3.2      farver_2.1.0
##  [43] pkgconfig_2.0.3     dbplyr_2.1.1        utf8_1.2.2
##  [46] labeling_0.4.2      tidyselect_1.1.1    rlang_0.4.12
##  [49] munsell_0.5.0       cellranger_1.1.0    tools_4.1.2
##  [52] cli_3.1.0           generics_0.1.1      evaluate_0.14
##  [55] fastmap_1.1.0       yaml_2.2.1          sys_3.4
##  [58] processx_3.5.2      knitr_1.37          fs_1.5.2
##  [61] randomForest_4.6-14 pbapply_1.5-0       slam_0.1-49
##  [64] ROI_1.0-0           xml2_1.3.3          rstudioapi_0.13
##  [67] compiler_4.1.2      e1071_1.7-9         reprex_2.0.1
##  [70] tweenr_1.0.2        stringi_1.7.6       highr_0.9
##  [73] ps_1.6.0            rgeos_0.5-9         Matrix_1.4-0
##  [76] classInt_0.4-3      vctrs_0.3.8         pillar_1.6.4
##  [79] lifecycle_1.0.1     BiocManager_1.30.16 data.table_1.14.2
##  [82] raster_3.5-11       R6_2.5.1            bookdown_0.24
##  [85] renv_0.14.0-148     KernSmooth_2.23-20  parallelly_1.30.0
##  [88] codetools_0.2-18    boot_1.3-28         rcdd_1.5
##  [91] MASS_7.3-54         rprojroot_2.0.2     withr_2.4.3
##  [94] mgcv_1.8-38         parallel_4.1.2      doSNOW_1.0.19
##  [97] hms_1.1.1           terra_1.4-22        quadprog_1.5-8
## [100] grid_4.1.2          class_7.3-19        minqa_1.2.4
## [103] rmarkdown_2.11      snakecase_0.11.0    err_0.2.0
## [106] itertools_0.1-3     numDeriv_2016.8-1.1 lubridate_1.8.0
```
